# Mechanical compressive forces increase PI3K output signaling in breast and pancreatic cancer cells

**DOI:** 10.1101/2021.10.18.464825

**Authors:** M. Di-Luoffo, C. Schmitter, E.C. Barrere, N. Therville, M. Chaouki, R. D’Angelo, S. Arcucci, B. Thibault, M. Delarue, J. Guillermet-Guibert

## Abstract

**Context:** Mechanical stresses, including compression, arise during cancer progression. In solid cancer, especially breast and pancreatic cancers, the rapid tumor growth and the environment remodeling explain their high intensity of compressive forces. However, the sensitivity of compressed cells to targeted therapies remains poorly known.

**Results:** In breast and pancreatic cancer cells, pharmacological PI3K inactivation decreased cell number and induced apoptosis. These effects were accentuated when we applied 2D compression forces in mechanically responsive cells. Compression selectively induced overexpression of PI3K isoforms and PI3K/AKT pathway activation. Further, transcriptional effects of PI3K inhibition and compression converged to control the expression of an autophagy regulator, GABARAP, which level was inversely associated with PI3K inhibitor sensitivity under compression. Compression alone blocked autophagy flux in all tested cells, while inactivation of basal PI3K activity restored autophagy flux only in mechanically non-responsive compressed cells.

**Conclusion:** This study provides direct evidence for the role of PI3K/AKT pathway in compression-induced mechanotransduction. PI3K inhibition promotes apoptosis or autophagy, explaining PI3K importance to control cancer cell survival under compression.

## 1. Introduction

In tissues, all cells are subjected to mechanical stresses, which correspond to the force per unit surface exerting onto the cell surface (measured in N/m^2^ or Pascal, Pa), that can notably be transmitted to the nucleus (Lomakin AJ et al, 2020). Physically, the applied force can be either normal or tangential to the surface, causing the cell to deform according to its material properties. Cells encounter three types of mechanical stresses: shear, tensile and compressive stress (Northcott JM et al, 2018). These mechanical interactions emerge from cell-cell or cell-substrate interaction (Levental KR et al, 2009). In cancers, compressive stress is poorly studied and its impact on cell proliferation and migration could depend on the magnitude, duration, and direction of applied forces, and the association to extra-cellular tissue components (Northcott JM et al, 2018). At homeostasis, epithelial cells sense compressive forces that limit their proliferation (Li Y et al, 2021); hence, the values of compressive forces applied to epithelial cells are expected to be low compared to pathological condition. During solid tumor development, tumor cells proliferate rapidly, and this situation is associated with an increase of mechanical forces and increased of internal rigidity (Therville N et al, 2019). In tumors, internal rigidity varies from 10 to 50 kPa (Therville N et al, 2019); however the magnitude of interstitial pressure close to tumor cells is of the order of the hundred of Pa (Provenzano PP et al, 2012). Rigidity of the stroma in which the tumor cells proliferate is increasing, promoting tumor cell confinement (Therville N et al, 2019). Those events lead to the emergence of large compressive stresses applied to confined tumor cells. *In vitro* 3D models of solid cancer cell growth under confinement show a cell proliferation decrease (Delarue M et al, 2014), while colon cancer cell compression without confinement promotes cancer cell growth (Mary G et al, 2022). Moreover, compression can increase cancer cell invasive capabilities (Kalli M et al, 2022, Kalli M et al, 2019a). Confinement induces compression stress and increases cell resistance to chemotherapeutic treatments; the latter is notably due to a maintained cell survival and reduced proliferation, limiting the effect of both cytotoxic/cytostatic agents (Delarue M et al, 2014, Rizzuti IF et al, 2020). Currently, there is no way to predict what will be the cellular physiological response of compression in cancer cells.

Once exerted on a cell, a mechanical stress induces a mechanotransduction response, coupled to modification of gene expression, which is largely associated with Hippo pathway activation in response to tensile stress (Cobbaut M et al, 2020). In tumors, Hippo pathway containing transcriptional regulators YAP/TAZ can reprogram cancer cells into cancer stem cells and incite tumor initiation, progression, and metastasis (Cobbaut M et al, 2020). Further, the Hippo pathway crosstalks with morphogenetic signals, such as Wnt growth factors, and is also regulated by Rho and G protein-coupled receptor (GPCR), cAMP and PKA pathways (Piccolo S et al, 2014). In the last decade, research has highlighted the interconnections of signaling pathways, and other key intracellular signals involved in mechanotransduction were identified (Di-Luoffo M et al, 2021, Schmitter C et al, 2023). In this context, the pivotal role of phosphoinositide 3-kinases (PI3Ks) in mechanotransduction of cancers is emerging (Borreguero-Munoz N et al, 2019, Zhao Y et al, 2018). However, PI3K pathway regulation under compressive stress response is not completely elucidated.

PI3K proteins can be divided into three classes (I–III)(Vanhaesebroeck B et al, 2010). In the literature, the term PI3K refers usually to class I PI3K, which consists in one p110 catalytic subunit and one p85 regulatory subunit excepted for PI3Kγ which harbored p87/p101 regulatory subunits. This class, composed of four enzymes (α, β, δ, γ), with non-redundant functions (Arcucci S et al, 2021, Vanhaesebroeck B et al, 2010) and non-redundant roles in cancer (Pons-Tostivint E et al, 2017), generates phosphatidylinositol 3,4,5-trisphosphate (PtdIns-3,4,5-P3 or PIP3) from phosphatidylinositol 4,5-bisphosphate (PtdIns-4,5-P2 or PIP2) (Vanhaesebroeck B et al, 2010). In pancreatic cancer cells, compression-induced PI3K activation promotes migratory phenotype involving an autocrine auto-stimulation loop (Kalli M et al, 2022, Kalli M et al, 2019a, Thibault B et al, 2021). Class I PI3Ks are upstream activators of YAP/TAZ transcriptional pathway under tensile stress, positioning class I PI3Ks proteins as potential regulators of an essential mechanotransduction signaling (Zhao Y et al, 2018). In breast cancer cells, *in vivo* overexpression of PI3Kβ sensitizes untransformed cells to YAP/TAZ-induced oncogenicity (Zhao Y et al, 2018). Besides, PI3K/AKT signaling closely regulates autophagy process (Heras-Sandoval D et al, 2014, Xu Z et al, 2020). Further, autophagy process was known to be induced by mechanical shear and tensile stresses in cells in pathological conditions, and in the integration of physical constraints (Claude-Taupin A et al, 2021, King JS, 2012, King JS et al, 2011). Importantly, PI3K pathway is one of the most common aberrantly activated pathways in cancers. Genetic alterations of *PIK3CA* (encoding PI3Kα) lead to constitutively active PI3K/AKT pathway; constitutive PI3K/AKT activation is associated with poor prognosis in pancreatic cancer (Thibault B et al, 2021). Cancer dependency to PI3K activation (including breast and pancreatic cancers) provided the rationale for development of inhibitors targeting PI3K/AKT pathway (reviewed in (Ellis H & Ma CX, 2019)) for their treatment.

While the implication of PI3K under tensile stress is now recognized as affecting tumor cell fate (Stylianopoulos T, 2017), its role under compressive stress still remains elusive. In particular, the mechanotransduction following a compressive stress remains poorly characterized. Recent evidence showed that, amongst all pro-tumoral signaling pathways, class I PI3Ks appear to be critically involved in the adaptive response to compression (Kalli M et al, 2019a, Nam S et al, 2019) (reviewed in (Di-Luoffo M et al, 2021)). Hence, PI3K targeting small molecule inhibitors might be relevant therapies to annihilate this adaptive response and lead to cancer cell death. To test this hypothesis, we thus determined the importance of class I PI3K activity in cancer cell lines known to be sensitive to PI3K inhibitors and, using *in silico* transcriptome analysis and mRNA/protein validation, we identified GABARAP as a common molecular determinant associated with autophagy controlled by compression forces. Breast and pancreatic cancer cells were studied here because mammary and pancreatic solid cancers are commonly subjected to mechanical stress during their development. The purpose of this study is to provide proof-of-concept showing that the therapeutic impact of targeted therapies is dependent on compressive mechanical context.

## 2. Results

### 2.1 PI3K pathway inhibition decreases cell number and increases apoptotic cell death in breast and pancreatic cancer cells

First, breast (MCF-7 and MDA-MB-231) and pancreatic (CAPAN-1 and PANC-1) cancer cell lines were treated using a class I pan-PI3K inhibitor (GDC-0941 (10 µM)) or vehicle (Figure 1A).

**Figure 1.**
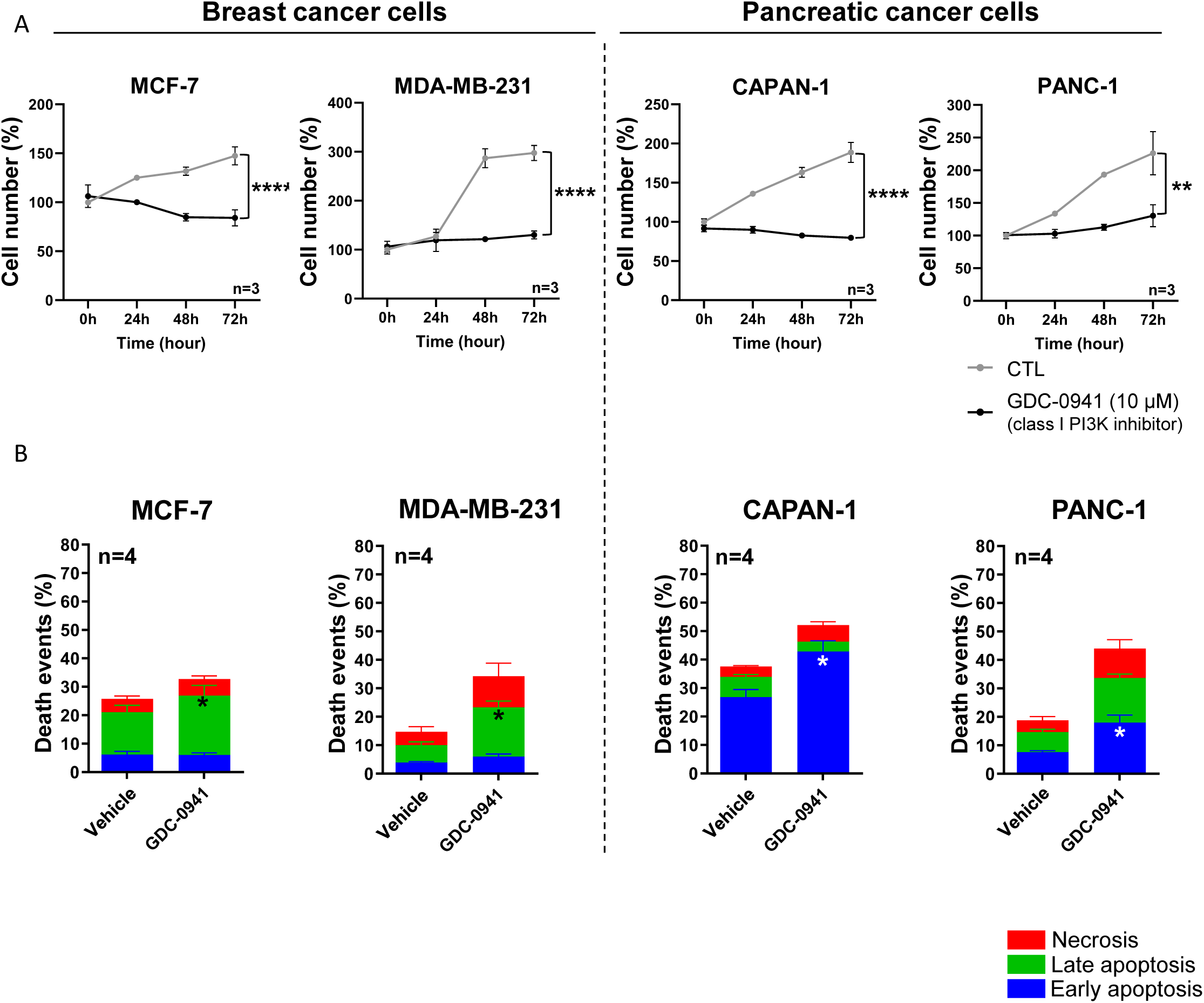
Cell number and death events in MCF-7/MDA-MB-231 breast and CAPAN-1/PANC-1 pancreatic cancer cells after inhibition of class I PI3K. After cell treatment, normalized cell number was quantified using Crystal Violet staining (**A-B**). **A.** Cell number was analyzed at 0, 24, 48 and 72 hours in MCF-7, MDA-MB-231, CAPAN-1 and PANC-1 cells, treated with class I pan-PI3K inhibitor (GDC-0941 (10µM)) compared to vehicle (CTL). **B.** After 24h pan-PI3K inhibitor treatment, cells were stained with a FITC-Annexin V/PI apoptosis detection kit. FITC-Annexin staining and Propidium Iodide (PI) incorporation were measured in cells using a flow cytometer and analyzed using FlowJo software. Early/late apoptotic and necrotic cell populations were measured in MCF-7, MDA-MB-231, CAPAN-1 and PANC-1 cells after class I pan-PI3K inhibitor (GDC-0941 (10µM)) treatment compared to vehicle (CTL). Detailed results of living cells, early/late apoptosis and necrosis were presented in Figure S4. nu=normalized unit. Results are presented as mean, +/- SEM, n=4. p-value after two-way ANOVA; *p-value<0.05.

Interestingly, 72h treatment with GDC-0941 (10 µM) significantly decreased the crystal violet staining indicative of cell number (cell number for short) in both breast and pancreatic cell lines compared to vehicle (Figure 1A, S1) which shows these cells as sensitive to the pan-PI3K inhibitor. These results were confirmed in two additional breast MDA-MB-468 and pancreatic Mia-Paca-2 cell lines (Figure S2A). To investigate the cell death mechanism involved in cell number decrease after PI3K pharmacological treatment, we performed Annexin V/Propidium Iodide (PI) cytometry experiments (Figure 1B, S2). The basal cell death status was different in each cell line. Interestingly, GDC-0941 significantly increased early or late apoptosis in pancreatic cancer cells (PANC-1, CAPAN-1; Figure 1B, S3) or in breast cancer cells (MCF-7, MDA-MB-231; Figure 1B, S2), respectively.

These data corroborate that class I PI3K (PI3Kα, β, δ, γ) are necessary for breast (MCF-7, MDA-MB-231) and pancreatic (PANC-1 and CAPAN-1) cancer cell survival.

### 2.2 PI3K pathway inhibition decreases cell number and increases apoptotic cell death under compression in mechanosensitive breast and pancreatic cancer cells

To investigate the impact of the PI3K/AKT pathway in mechanotransduction following compressive stress, we developed a compression device to compress cells in 2D (Figure 2). The breast and pancreatic cells were compressed with a calibrated 200Pa pressure per cell adjusting the total mass with small metal weights on agarose pad (Figure 2). The controls were performed adding the equivalent volume of the agarose pad medium. We applied both pharmacological inhibition (GDC-0941 (10 µM)) and a 200 Pa 2D compressive forces at single cell level. Next, we quantified cell number after 0, 24h, 48h and 72h compression. Interestingly, 200 Pa compression associated with pan-PI3K inhibitor (200 Pa + GDC-0941) significantly decreased cell number in (MCF-7 and CAPAN-1) now identified as mechanically responsive cells, while GDC-0941 did not affect cell number in mechanically non-responsive cells (MDA-MB-231 and PANC-1) (Figure 3A). These results were confirmed in an additional mechanically responsive Mia-Paca-2 cell line and a mechanically non-responsive MDA-MB-468 cells (Figure S2B).

**Figure 2.**
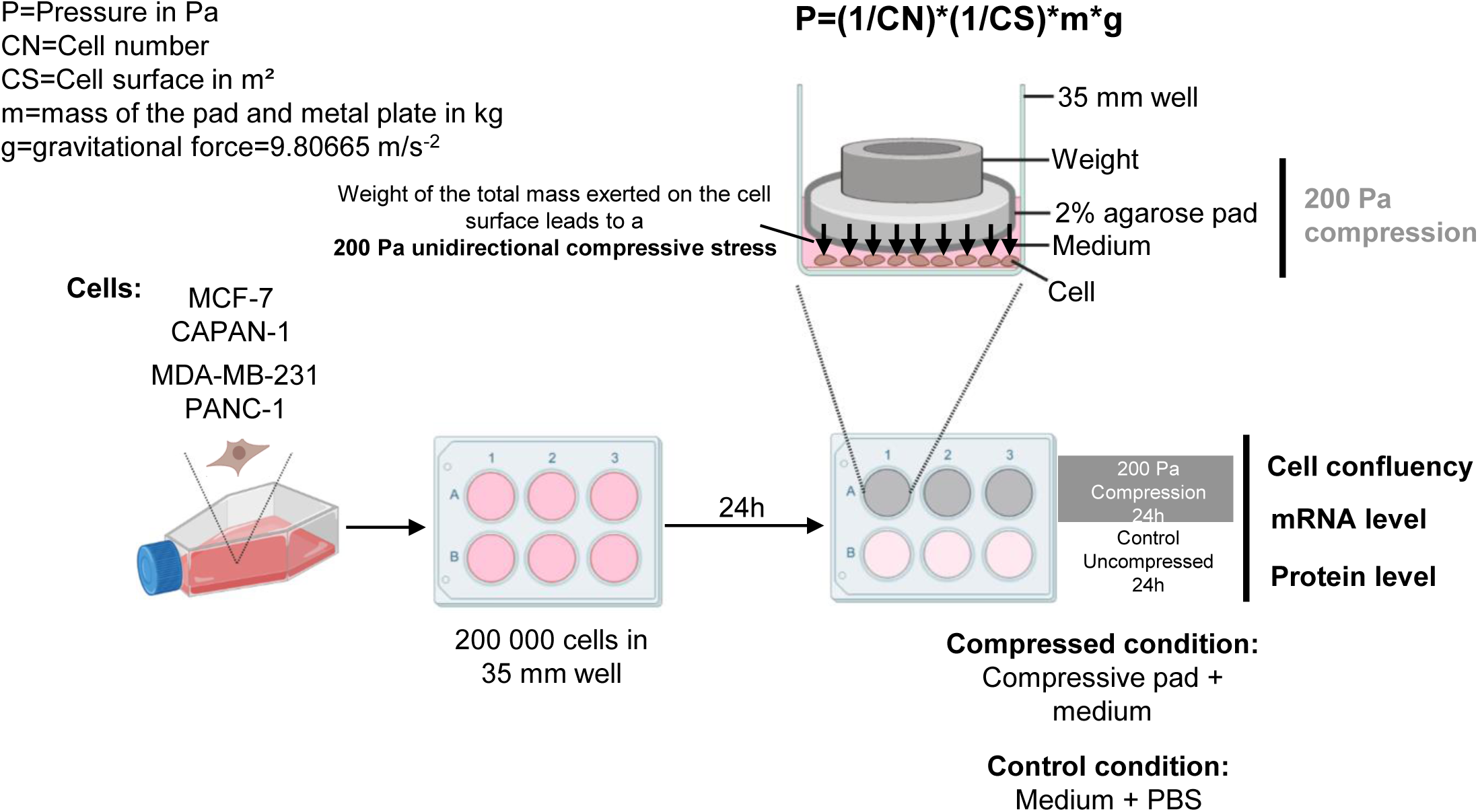
2D compression protocol applied in MCF-7, CAPAN-1, MDA-MB-231 and PANC-1 cancer cells. MCF-7, CAPAN-1, MDA-MB-231 and PANC-1 cells were plated in a 35 mm petri dish. 24h after plating, a 2% low gelling temperature agarose pad in Phosphate Buffered Saline (PBS) was deposited on cells. 2D compression was calibrated at 200 Pa per cell adjusting the mass with small weight on 2% agarose pad and calculated using the formula: P=(1/CN)*(1/CS)*m*g (with P=Pressure in Pa; CN=Cell number; CS=Cell surface in m²; m=mass of the pad and metal plate in kg; g=gravitational force=9.80665 m/s-2). Controls were performed adding the volume of PBS equivalent to the volume of agarose pad. After 24h of compression, cells were harvested, and cell number was calculated and RNA/proteins levels were quantified.

**Figure 3.**
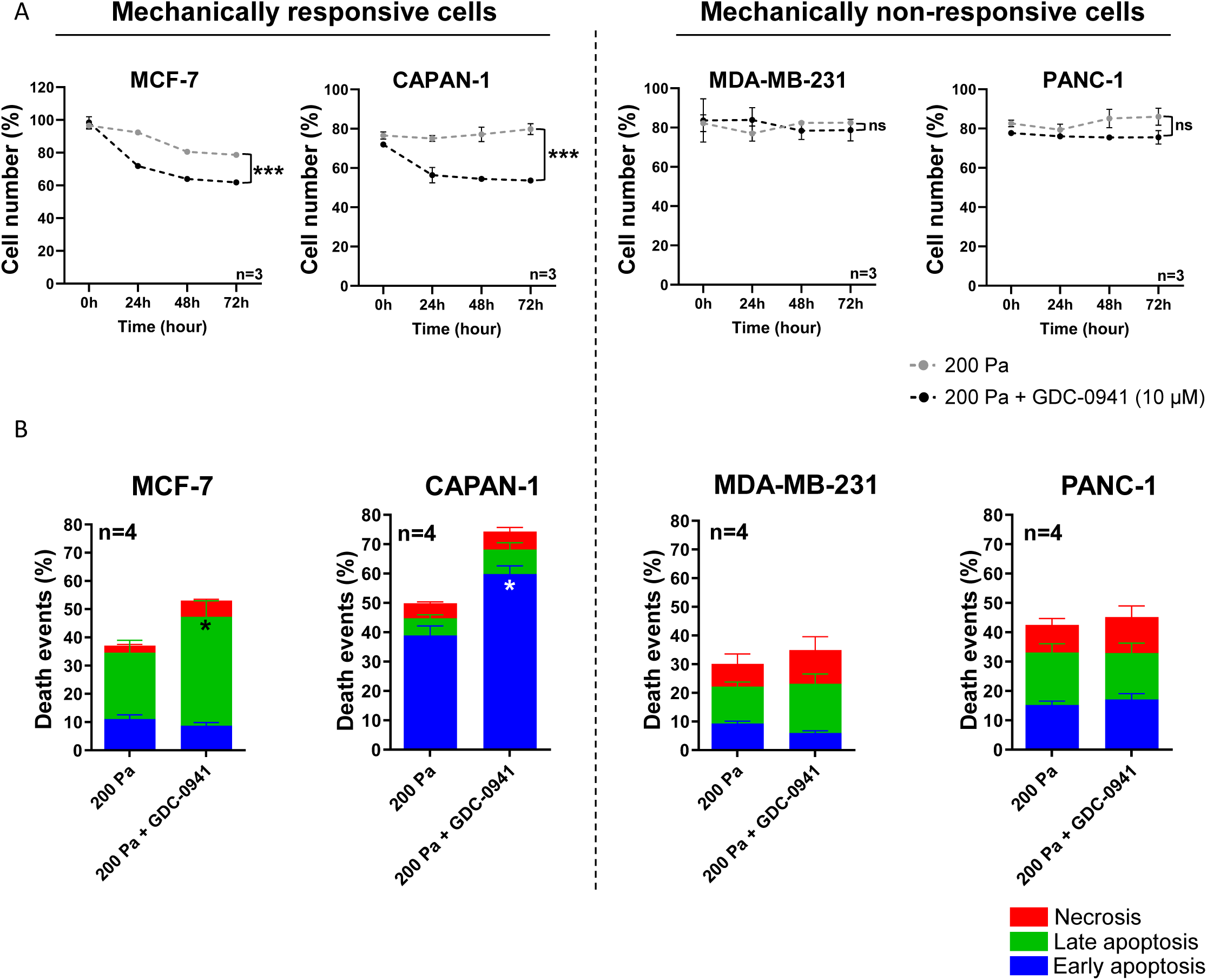
Cell number and death events in MCF-7/CAPAN-1 mechanically responsive and PANC-1/MDA-MB-231 non-responsive cells associated with inhibition of class I PI3K. **A.** Normalized cell number was analyzed at 0, 24, 48 and 72 hours in MCF-7, CAPAN-1, MDA-MB-231 and PANC-1 cells after 200 Pa compression (200 Pa) and class I pan-PI3K inhibitor (200 Pa + GDC-0941 (10µM)) compared to 200 Pa compressed cells (200 Pa). After cell treatment, cell number was quantified using Crystal Violet staining. **B.** After 24h pan-PI3K inhibitor treatment, cells were stained with a FITC-Annexin V/PI apoptosis detection kit. FITC-Annexin staining and Propidium Iodide (PI) incorporation were measured in cells using a flow cytometer and analyzed using FlowJo software. Early/late apoptotic and necrotic cell populations were measured in in MCF-7, CAPAN-1, MDA-MB-231 and PANC-1 cells after 200 Pa compression and class I pan-PI3K inhibitor (200 Pa + GDC-0941 (10µM)) treatment compared to compressed cells (200 Pa). Detailed results of living cells, early/late apoptosis and necrosis were presented in Figure S4. nu=normalized unit. Results are presented as mean, +/- SEM, n=4. p-value after two-way ANOVA; *p-value<0.05.

Moreover, 200 Pa compression associated with pan-PI3K inhibitor (200 Pa + GDC-0941) significantly increased late apoptosis in MCF-7 breast cancer cells and early apoptosis in CAPAN-1 pancreatic cells that are mechanically responsive cells. However, 200 Pa + GDC-0941 did not affect apoptosis process in mechanically non-responsive cells (MDA-MB-231 and PANC-1) (Figure 3B).

These data support that the cell number decrease in mechanically responsive cells comes from an increase in early and late apoptosis process, which was not affected in mechanically non-responsive cells.

### 2.3 Compression activates PI3K pathway in mechanically responsive cells

To experimentally validate the compression impact on PI3K pathway, we next compressed mechanically responsive cells (MCF-7 and CAPAN-1) and mechanically non-responsive cells (MDA-MB-231 and PANC-1). 200 Pa compression significantly increased by 3.5-fold and 2.7-fold (p-value=0.04) p-AKT^S473^/total AKT ratio in MCF-7 and CAPAN-1 cells, indicative of PI3K activation in mechanically responsive cells. Further, we analyzed gene expression of the four class I PI3K genes (*PIK3CA*, *PIK3CB*, *PIK3CD* and *PIK3CG*). MCF-7 cells harbored a 2.9-fold and 6.2-fold significant overexpression of *PIK3CA* and *PIK3CD* and CAPAN-1 cells harbored a 2.7-fold (p-value=0.04), 1.5-fold (p-value=0.02) and 4.9-fold (p-value=0.03) significant overexpression of *PIK3CB, PIK3CD* and *PIK3CG* respectively (Figure S4). Alternatively, 200 Pa compression did not significantly modulate p-AKT^S473^/total AKT ratio in mechanically non-responsive cells MDA-MB-231 and PANC-1 (p-value=0.21 and 0.26 respectively) (Figure 4). Only *PIK3CG* expression was 11.7-fold significantly increased in PANC-1 (p-value=0.02) (Figure S4). These results were confirmed in two additional breast MDA-MB-468 and pancreatic Mia-Paca-2 cell lines (Figure S2C). These experimental results show that the activation of PI3K pathway as assessed by AKT phosphorylation under 200 Pa compression is mostly increased in mechanically responsive cells (MCF-7 and CAPAN-1) while was not affected in mechanically non-responsive cells (MDA-MB-231 and PANC-1). In addition, *PIK3CA*, *PIK3CB*, *PIK3CD* and *PIK3CG* mRNA overexpression in mechanically responsive cells confirm the AKT activation passes through important PI3K pathway members such as class I PI3K (PI3Kα, β, δ, γ).

**Figure 4.**
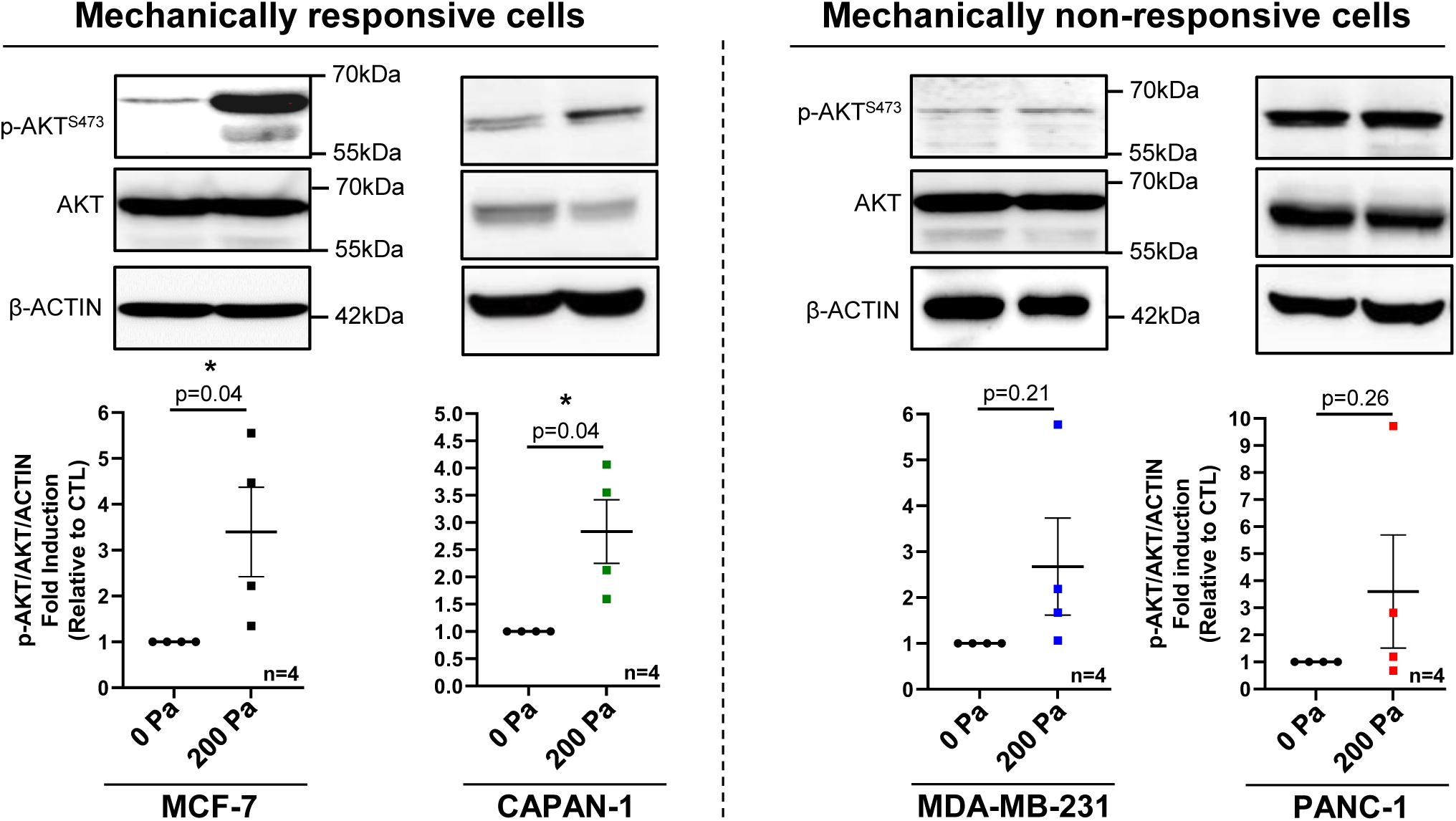
PI3K signaling in MCF-7/CAPAN-1 mechanically responsive and PANC-1/MDA-MB-231 non-responsive cells. Representative western blots of p-AKT^S473^ and AKT in mechanically responsive cells (MCF-7/CAPAN-1) and in mechanically non-responsive cells (PANC-1/MDA-MB-231) after 200 Pa compression (200 Pa) and non-compressed cells (0 Pa) for 24h. β-ACTIN was used as loading control. p-AKT^S473^/AKT quantitative analyses were performed using ImageJ software. Results are presented as mean, +/- SEM, n=4. *p-value<0.05. Representation of *PIK3CA*, *PIK3CB*, *PIK3CD*, *PIK3CG* mRNA expressions in mechanically responsive cells (left panel) and in mechanically non-responsive cells (right panel) after 200 Pa compression were presented in Figure S5.

### 2.4 PI3K-AKT pathway-regulated gene expression is sensitive to compression in breast cancer cells

To dive deeper into the molecular mechanism, we decided to analyze transcriptomics data and to understand how compressive stress altered gene signature of cell signaling pathways, including PI3K pathway. To this end, we searched for datasets obtained upon compression of cancer cells in Gene Expression Omnibus database (GEO), where public sequencing data from published studies are available (Figure 5A). In GEO database, one transcriptomic dataset obtained under cell compression is currently available. Kim *et al*. 2019 (Kim BG et al, 2019) compressed breast cancer cells using compressive pads applied on top of 2D cell layers. In this study, Kim *et al*. focused on and analyzed the action of compression on stromal cells. They observed that mechanical stress in these cells promoted a specific metabolic gene signature increasing glycolysis and lactate production (Cairns RA et al, 2011). Here, we compared the gene expression evolution of the two breast cancer cell lines MDA-MB-231 and MCF-7 that harbor decreased cell number under compression. Both breast cancer cell lines were compressed by a global compression applied on all cells ranging from 0 to 8kPa, and gene expression profile was analyzed. We first searched for gene signature enrichment analysis using canonical signatures from Gene Set Enrichment Analysis (GSEA) (Figure 5A). The gene signatures in different pathways were normalized into transcripts per million and compared to housekeeping gene expression such as *ACTINB, LAMINA1* and *LAMIN* gene class. The expressions of these housekeeping genes were not affected by increasing compression (Figure S5A). Under increasing compression [0], [0.3-0.8], [1.5-4.0], [7.0-8.0] kPa, PI3K-AKT signature gene expression was significantly overexpressed in MDA-MB-231 cells, suggesting a global overexpression of genes belonging to these pathways, while it was found already high in MCF-7 cells (Figure S5B, left panel). Those results are in line with known genetic status of MCF-7 cells, in which PI3Kα is constitutively active by mutation of its encoding gene (*PIK3CA*^E545K^). However, *PIK3CA*, *PIK3CB* and *PIK3CD* expressions increased in MCF-7 cells and remained stable in MDA-MB-231 cell line (Figure S5B, right panel).

**Figure 5.**
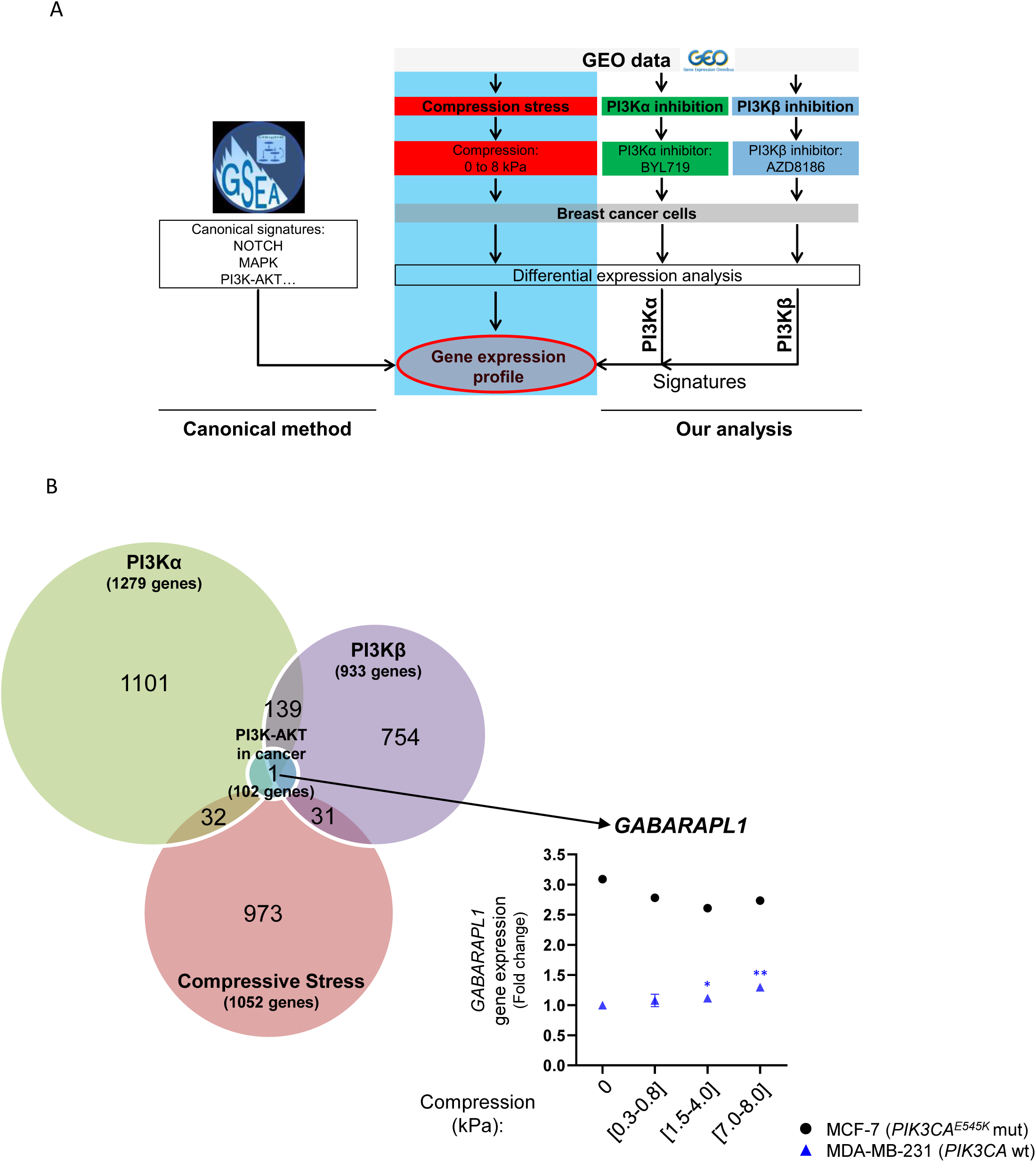
Comparative analysis between compressive stress, PI3Kα/PI3Kβ and PI3K-AKT signaling in cancer gene signatures. **A.** Workflow analysis comparing the lists of 1279, 933 and 1052 differentially expressed genes in PI3Kα signature, PI3Kβ signature, compressive stress signature and 102 genes of REACTOME PI3K-AKT signaling in cancer signature were compared. **B.** Venn diagram shows the overlapping between PI3Kα signature, PI3Kβ signature, compressive stress and REACTOME PI3K-AKT signaling in cancer. Overlapping between compressive stress/PI3Kα signature (32 genes), compressive stress/PI3Kβ signature (31 genes) and PI3Kα/PI3Kβ signatures (139 genes). The only common gene differentially expressed in PI3Kα inhibition/PI3Kβinhibition/compressive stress was *GABARAPL1*. MDA-MB-231 (*PIK3CA^wt^*; blue triangles) and MCF-7 (PIK3CA^E545K^ mutated, black dots) breast cancer cells were gradually compressed (0 kPa; [0.3 to 0.8 kPa]; [1.5 to 4.0 kPa]; [7.0 to 8.0 kPa] respectively) and *GABARAPL1* gene expression was quantified using Agilent microarray (Data available in https://www.ncbi.nlm.nih.gov/geo/query/acc.cgi?acc=GSE133134).

Interestingly, YAP/TAZ pathway gene signature was not significantly affected by increasing compression in both MCF-7 and MDA-MB-231 cells (Figure S5C, left panel). These transcriptomic results were experimentally confirmed by RT-qPCR analyzing gene expression of YAP/TAZ pathway members such as *TEAD1* and *c-JUN* and of downstream effectors of YAP/TAZ pathway such as *CCN1* and *CCN2.* Only the *CCN2* mRNA expression was 1.36-fold significantly increased of in MCF-7 mechanically responsive cells (Figure S6A). However, the expression level of other genes was not significantly modulated using 200 Pa compression (200 Pa) in mechanically responsive cells (MCF-7 and CAPAN-1) as well as in mechanically non-responsive cells (PANC-1 and MDA-MB-231) (Figure S6A). Those transcriptomic data were confirmed by other molecular assays such as the study of YAP phosphorylation that control its nuclear translocation. The p-YAP^S127^/YAP ratio was not significantly changed under 200 Pa compression in all cell lines (Figure S6B).

Taken together, these data showed an increased gene expression of PI3K family members in cells growing under compression (from 0 to 8 kPa) especially in mechanically responsive MCF-7 breast cancer cells.

### 2.5 PI3K pathway-regulated and compression-regulated gene expression share common targets and control autophagy gene expression in breast cancer cells

To go further using transcriptomic analysis and investigate the specific role of PI3K isoforms in the response to compression, we analyzed data from two more studies, which used selective inhibitors of PI3K isoforms: Bosch *et al*., 2015 (Bosch A et al, 2015) and Lynch *et al*., 2017 (Lynch JT et al, 2017). The authors used BYL719 or AZD8186 compounds described as selective inhibitors of PI3Kα or PI3Kβ respectively, in mutant *PIK3CA* MCF-7 and mutant *PTEN* HCC70 breast cancer cells. Mutant *PTEN* leads to increased PI3K activity (Chalhoub N et al, 2009). We next used gene expression data from PI3Kα inhibition in MCF-7 and PI3Kβ inhibition in HCC70 cells to define PI3Kα and PI3Kβ signatures. Selective transcriptional targets were crossed compared with the compression-specific transcriptional targets that we previously identified (Figure 5A).

After PI3Kα inhibition in MCF-7, 1279 genes had a significantly altered expression. After PI3Kβ inhibition in HCC70 cells, gene expression of 933 genes was also significantly affected. Gene expression of 1052 targets was affected in compressive conditions ([0.3-0.8], [1.5-4.0] and [7.0-8.0] kPa compared to [0] kPa) in MCF-7 cells. We compared these signatures to the list of the 102 genes found in canonical “PI3K-AKT REACTOME signaling pathway in cancer” signature (Figure 5B). PI3Kα and PI3Kβ signatures overlapped with the regulation of 139 genes. Compressive stress gene signature mostly overlapped either with PI3Kα or PI3Kβ signatures, on non-common 32 and 31 genes, suggesting a differential effect on isoform activation in response to compression.

Finally, GABA type A receptor associated protein like 1 gene (*GABARAPL1)*, coding for GABARAP structural protein of the autophagosome, was the only shared target gene between PI3Kα signature, PI3Kβ signature, compressive stress signature and “PI3K-AKT REACTOME signaling pathway in cancer” signature (Figure 5B). *GABARAPL1* and autophagy GSEA pathway gene signatures were affected under increasing compression in breast cancer cells (Figures 5B, S5C). In details, *GABARAPL1* gene expression was significantly upregulated after PI3Kα (x1.90; p-value=0.08) and PI3Kβ (x1.42; p-value=0.006) inhibitions. Under increasing compression from [0] kPa up to [7.0-8.0] kPa, *GABARAPL1* gene expression was significantly increased in mechanically non-responsive MDA-MB-231 cells, but not in mechanically responsive MCF-7 cells (Figure 5B, right panel). Moreover, under increasing compression, the canonical autophagy GSEA pathway gene expression significantly increased in MDA-MB-231 cells and decreased in constitutive PI3K activated MCF-7 cells (Figure S5C, right panel). These analyses suggest an association between PI3K activation and an increase of autophagy process involved in the non-response of cells to mechanical compressive forces. Moreover, the *GABARAPL1* expression alterations converge towards the regulation of autophagosome structural genes.

### 2.6 Compression modulates the expression of autophagosome structure proteins in breast and pancreatic cancer cells

Autophagy process was proposed as a therapeutic target for various tumors (Xu Z et al, 2020); inhibition of autophagy in cancer cells decreases their survival. PI3Kα and PI3Kβ are known to be positive and negative regulators of autophagy process (Heras-Sandoval D et al, 2014, Xu Z et al, 2020). When comparing PI3Kα and PI3Kβ signatures, PI3K-AKT REACTOME canonical pathway and compressive stress selective gene expression, only *GABARAPL1* gene was significantly affected and overlapped in these four signatures (Figure 5). Briefly, this gene encodes for a structural protein of the autophagosome and autophagolysosome involved in autophagy process (reviewed in (Le Grand JN et al, 2011)). To validate transcriptomics data, we decided to analyze autophagy as well as *GABARAPL1* gene expressions in compressed cells. *GABARAPL1* gene expression was significantly increased in mechanically non-responsive cells (Figure 6). More precisely, *GABARAPL1* gene expression was increased by 2.1-fold (p-value=0.01) in MDA-MB-231 accordingly to transcriptomic data analysis and by 3.9-fold (p-value=0.03) in PANC-1 cells (Figure 6A, right panel). These results were confirmed with GABARAP protein level which was 5.9-fold significantly increased in MDA-MB-231 (p-value=0.04) and remains high between uncompressed (0 Pa) and compressed (200 Pa) PANC-1 cells (Figure 6B, right panel). In contrast, GABARAP mRNA and protein levels were not significantly modulated in mechanically responsive cells, excepted for the decrease in GABARAP protein level in CAPAN-1 cells in 200 Pa compression condition (Figure 6A-B, left panel), suggesting a decrease of autophagy process in mechanically sensible cells. Further, in the mechanically responsive Mia-Paca-2 cell line, compression did not induce GABARAP protein overexpression.

**Figure 6.**
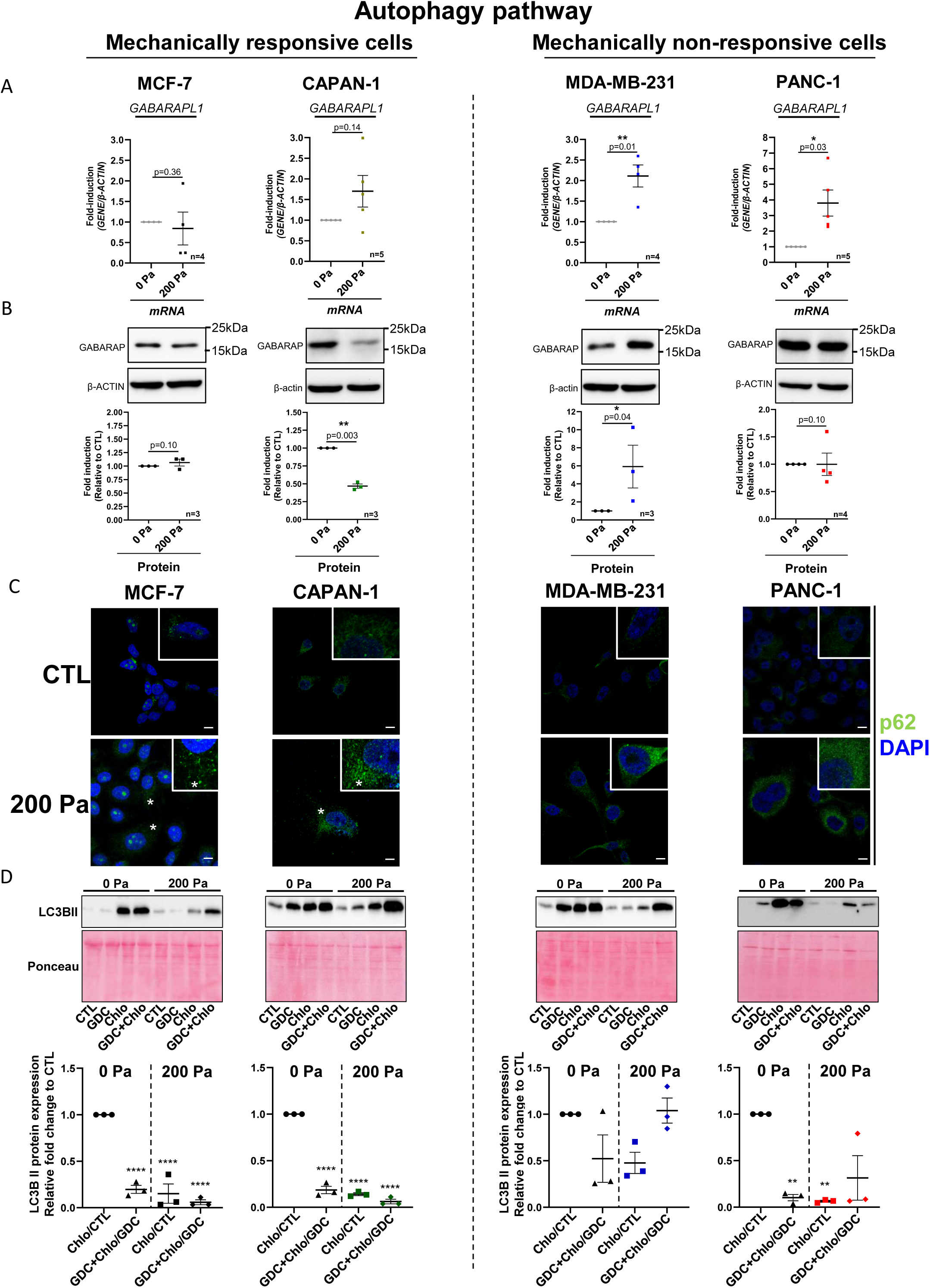
Autophagy flux in MCF-7/CAPAN-1 mechanically responsive and PANC-1/MDA-MB-231 non-responsive cells. **A.** *GABARAPL1* mRNA expressions in mechanically responsive cells (MCF-7/CAPAN-1) and in mechanically non-responsive cells (PANC-1/MDA-MB-231) after 200 Pa compression (200 Pa) and non-compressed cells (0 Pa) for 24h. β-ACTIN was used as housekeeping gene. Results are presented as mean, +/- SEM. n≥4 for each gene. p=p-value. *p-value<0.05; **p-value<0.01. **B**. Representative western blots of GABARAP in mechanically responsive cells (MCF-7/CAPAN-1) and in mechanically non-responsive cells (PANC-1/MDA-MB-231) after 200 Pa compression (200 Pa) and non-compressed cells (0 Pa) for 24h. β-ACTIN was used as loading control. GABARAP/β-ACTIN quantitative analyses were performed using ImageJ software. Results are presented as mean, +/- SEM, n≥3. p=p-value. *p-value<0.05; **p-value<0.01. **C.** p62/SQSTM1 staining in mechanically responsive cells (MCF-7/CAPAN-1) and in mechanically non-responsive cells (PANC-1/MDA-MB-231) after 200 Pa compression (200 Pa) and non-compressed cells (0 Pa) for 24h. Green fluorescence corresponds to p62/SQSTM1 and nuclear signal corresponds to DAPI. * represent puncti. Scale bar correspond to 10µm. n=3. **D.** Representative western blots of LC3B II in mechanically responsive cells (MCF-7/CAPAN-1) and in mechanically non-responsive cells (PANC-1/MDA-MB-231) after 200 Pa compression (200 Pa) and non-compressed cells (0 Pa) for 24h. The quantification represents the ratio between 10 µM chloroquine and control conditions (Chlo/CTL) and 10 µM GDC-0941 + 10 µM chloroquine to 10 µM GDC-0941 (GDC+Chlo/GDC) in 200 Pa compression (200 Pa) and uncompressed (0 Pa) conditions in all cells. Ponceau S was used as loading control. LC3B II/ Ponceau S quantitative analyses were performed using ImageJ software and normalized to CTL 0 Pa. Results are presented as mean, +/- SEM, n≥3. p-value after two-way ANOVA; **p-value<0.01; ****p-value<0.0001.

However, in the mechanically non-responsive MDA-MB-468 cells, 200 Pa compression prompted a GABARAP protein increased level (Figure S2B).

These experimental results show that GABARAP level remains low or decreases in mechanically responsive cells. However, GABARAP level increases or remains high in mechanically non-responsive cells after compression, cells which are mechanically non-responsive. The functional link between mechanical insensitivity, autophagy flux and inactivation of the PI3K pathway therefore will be studied in the remainder of this study.

### 2.7 Pharmaceutical inhibition of PI3K decreases the autophagy flux only in mechanically responsive cells

Then, we decided to investigate the cellular localization of the autophagosome cargo p62/SQSTM1 by immunofluorescence. Its accumulation in puncta is usually observed when autophagy process is blocked (Bjorkoy G et al, 2009). We observed the p62/SQSTM1 level and localization in breast and pancreatic cancer cells under compression. A 200 Pa compression increased the density and the perinuclear localization of p62/SQSTM1 puncta in four cell lines (Figure 6C), while larger p62 puncta were only observed in mechano-responsive cells (Figure 6C, left panel).

We next confirmed whether PI3K inactivation in mechanically responsive cells (MCF-7 and CAPAN-1) might further block autophagy process under compression and quantified the level of autophagy flux. At autophagosome initiation, LC3B-I is lipidated in LC3B-II (Heras-Sandoval D et al, 2014, Xu Z et al, 2020). Next, LC3B-II autophagosomes are degraded by fusion with lysosomes. To assess the level of LC3B-II-autophagic flux, we used chloroquine (10 µM) that inhibits autophagy by impairing autophagosome fusion with lysosomes (Mauthe M et al, 2018), to determine the amount of LC3B-II-autophagosomes that is at this time point degraded by lysosomes. Comparing LC3B-II level of chloroquine-treated to controls in the different conditions is thus a mean to assess changes in LC3B-II-mediated autophagy flux. We next measured the chloroquine-induced LC3B-II levels in each condition (condition+Chlo/condition ratio) and compared them to the control ratio (Chlo/CTL). In mechanically responsive cells, the LC3B-II-autophagy flux was decreased in 0 Pa compression, upon PI3K inhibitor treatment, treating the MCF-7 and CAPAN-1 cell lines with pan-PI3K inhibitor (GDC-0941 (10 µM)) (GDC+Chlo/GDC) (Figure 6D, left panels). Further, 200 Pa compression condition decreased even more LC3B-II level with or without pan-PI3K inhibitor (Chlo/CTL or GDC+Chlo/GDC) (Figure 6D, left panels). However, in the mechanically non-responsive MDA-MB-231 cells, pan-PI3K inhibitor (GDC+Chlo/GDC) only or associated with 200 Pa compression did not affect significantly LC3B-II-autophagy flux (Figure 6D, right panels). In the mechanically non-responsive PANC-1 cells, pan-PI3K inhibitor (GDC+Chlo/GDC) decreased LC3B-II-autophagy flux without compression, however 200 Pa compression + pan-PI3K inhibitor (GDC+Chlo/GDC) did not affect LC3B-II-autophagy flux (Figure 6D, right panels). Taken together, these results showed that, while compression blocks autophagy, PI3K inactivation restores autophagy flux in mechanically non-responsive compressed cells.

## 3. Discussion

The impact of mechanical compressive forces on biological processes in cancer cells is still underexplored. Here, our major finding is that high intensity and static compression influences growth rate, cell death and autophagy process in breast and pancreatic cancer cells that are mechanically responsive or non-responsive cells. PI3K inhibition accentuates this effect, opening avenues for compression in combination with PI3K inhibitors as a therapeutic intervention (Figure 7).

**Figure 7.**
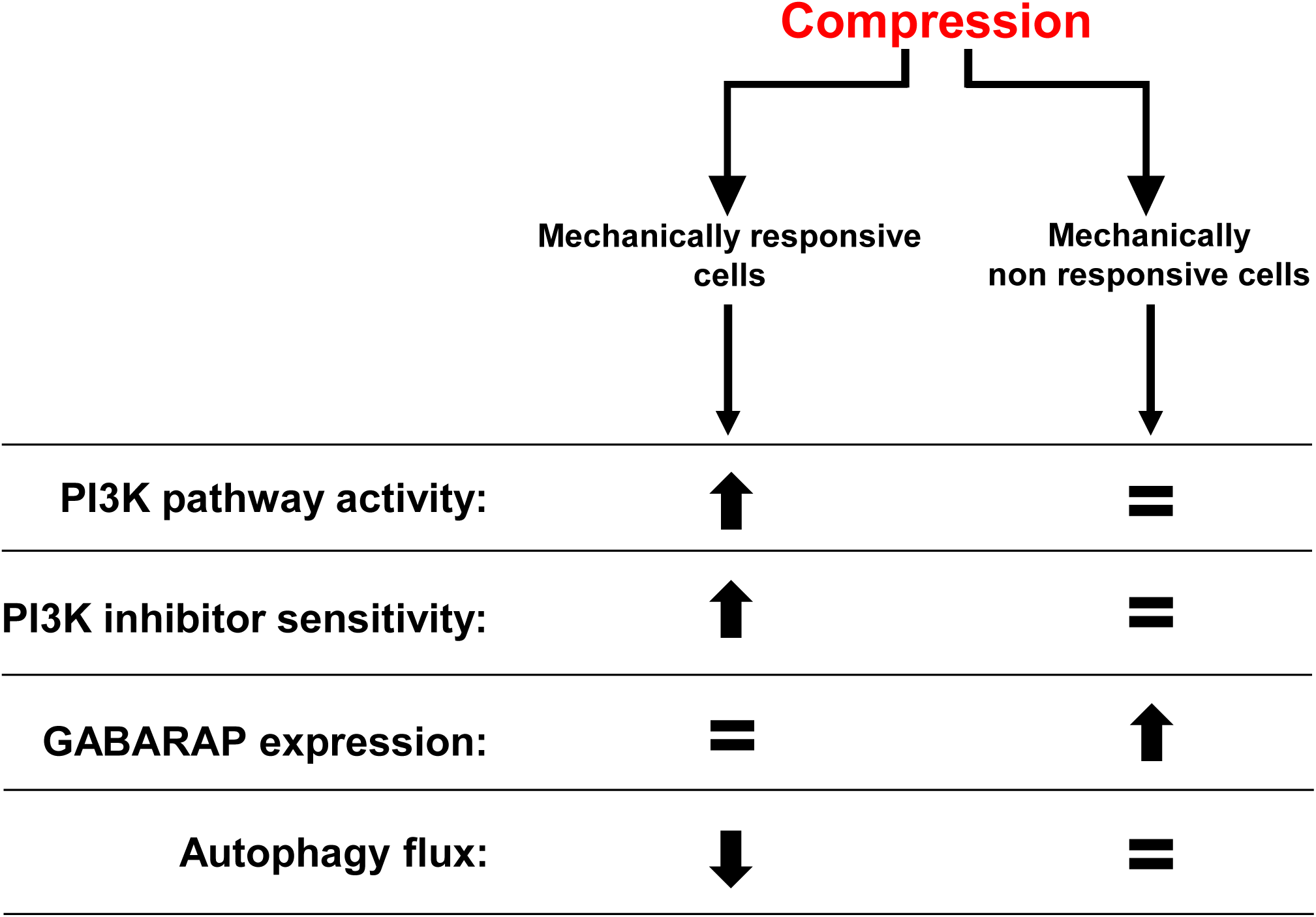
Balanced impact of compressive stress in mechanically responsive and non-responsive cells. PI3K pathway activity, PI3K inhibitor sensitivity, GABARAP expression and autophagy flux were modulated in mechanically responsive and non-responsive cells. The balance leans towards biological process over/down activation or actor over/down expression (black arrows) depending on the cell compressive status.

In terms of molecular mechanisms, the AKT activation by PI3K pathway has a well-described important role in survival signaling. Activated AKT phosphorylates and inhibits the pro-apoptotic BCL-2 family members BAD, BAX, CASPASE-9, GSK-3, and FOXO1, which tightly regulates apoptosis (reviewed in (Miricescu D et al, 2020)). Hence, besides the pro-migratory (Kalli M et al, 2019a) and proliferative (Nam S et al, 2019) effects of compression-induced PI3K activation, we show here in breast and pancreatic cancer cells that PI3K can also control cancer cell death processes (early or late apoptosis) under compression.

Through our unbiased approach, we also identify another cell process, autophagy, as critically involved in determining the biological output of compression in cancer cells. Autophagy is a cellular process that allows the orderly degradation and recycling of cellular components, hence providing a self-promoted nutrient source benefiting cell homeostasis and survival (Poillet-Perez L & White E, 2019). Autophagy is a highly conserved catabolic process, participating in the balance between cell proliferation and cell death in tumor and the tumor microenvironment regulation. This process is highly regulated by PI3K-AKT-mTOR pathway in various tumors. We know that inhibition of PI3K-AKT-mTOR induces autophagy as part of a tumor growth suppressor role in early cancerogenesis (Cui LH et al, 2019, Yang J et al, 2018). During cancer initiation, the autophagic flux participates to tissue homeostasis and helps to remove damaged cells. However, autophagy has a dual effect in cancer. In established cancers, autophagy promotes cancer cell survival and resistance to treatment. Autophagy is well known to be a key therapeutic strategy for a variety of tumors including pancreatic cancer (Bryant KL et al, 2019). Interestingly, PI3K activity controls autophagy in cancer cells with or without compression.

By comparing genes regulated by PI3Kα/PI3Kβ inhibitions, compressive stress and reactome PI3K-AKT signaling in cancer pathway lists of genes, we identified only *GABARAPL1* gene as shared gene deregulated in these three signatures. Experimental data confirmed the bioinformatics analysis. Compression altered GABARAP protein level as GABARAP protein level decreases or remains low in the two cell lines where PI3K inhibition further induces cell death under compression (mechanically responsive cells). GABARAP protein is known to be cleaved by ATG4B protease before its conjugation to phospholipids. This modified lipidated form is localized on the surface of autophagosomes and lysosomes, participating in their formation. It is therefore involved in autophagy process (reviewed in (Le Grand JN et al, 2011)). The differential expression of GABARAP protein raises the possibility of a modulation of autophagy flux (acceleration or blocking) and certainly of the stability of autophagosomes under compressive stress.

There are only limited studies that link induction of autophagy with mechanical stress (Claude-Taupin A et al, 2021, Xu Z et al, 2020). In *Dictyostelium discoideum*, compression activates autophagy in a mTORC1 independent manner (King JS et al, 2011). This mechanism, if confirmed in mammalian cells, could participate in cancer cell survival as well as resistance to chemotherapeutic treatments under compression. However, the fact that PI3K inhibition could revert or prevent the blockage of autophagy suggest that mammalian cells might have other mechanisms of cell adaptation to compression.

Under increasing compression, gene/protein expression or phosphorylation status of PI3K-AKT and autophagy pathway members were significantly modulated. However, and surprisingly, “YAP/TAZ *via* Hippo” or “YAP/TAZ Non Hippo” pathway gene signatures and expression of YAP/TAZ-regulated genes were not significantly affected in breast and pancreatic cancer cells under unidirectional 2D compressive stress (Figure S6). YAP/TAZ Hippo and Non Hippo pathways are known as key regulators of mechanotransduction under tensile stress (Totaro A et al, 2018). In compressive stress context, the implication of YAP/TAZ pathway was less investigated. It was assumed that this stress would induce the same signaling pathways as tensile stress. This assumption could have been also prompted by the fact that some techniques used to induce compression could also promote tensile stress (Chen Y et al, 2019, Kalli M et al, 2019b). It is possible that compression in our system could also create a tension on the edge of cell monolayer, that is known to be rapidly regulated (Venkova L et al, 2022); as we are studying the effect of compression at much higher time scales, the response to compression might prevail in our case. We could argue that a threshold of tensile stress may be needed to permit the activation of YAP/TAZ. If our data seem to confirm those findings in the context of unidirectional compressive stress, differential cytoskeleton remodeling cannot be completely ruled out in our experimental conditions. The exact type of cytoskeleton remodeling under each mechanical stress may also be key to understand cell mechanotransduction processes, as cells adapt to the environmental stresses. Finally, PI3K signal is permissive for tensile stress control of YAP/TAZ and targeting of PI3K could be a novel strategy to hinder the potential YAP/TAZ oncogenic dependence (Di-Luoffo M et al, 2021).

Altogether, we show that breast and pancreatic cancers are sensitive to compression (Kalli M et al, 2019a). In tumors, compression forces are mostly a consequence of tumor cell confinement in rigid microenvironment, that comprises huge ECM accumulation, deposits also called desmoplasia. These high compressive stress situations occur mainly in breast and pancreatic cancers (Thibault B et al, 2021, Verret B et al, 2019). We previously published that confinement reduced the efficiency of Gemcitabin and Paclitaxel in pancreatic cancer spheroids due to compression reduced proliferation rate (Rizzuti IF et al, 2020); (Schmitter C et al, 2023). Our current study provides an exploratory proof-of-concept of the impact of mechanical stresses on targetable oncogenic cell signaling. The efficiency of both signal-targeted therapies and chemotherapies combined with mechanotherapeutics should however be investigated in a wider range of solid cancers, where matrix remodeling is a key component of tumor progression. One widely studied strategy is to target activated PSC and CAFs which play an important role on ECM composition, stiffness and non-autonomous autophagy (Yang A et al, 2018); (Mukhopadhyay S et al, 2023); (Below CR et al, 2022); (Yang D et al, 2023). Further, ECM accumulation and desmoplasia may even be heterogeneously distributed in each pancreatic tumor and evolve during progressive stages of the disease (Provenzano PP et al, 2012); tumoral rigidity is also heterogeneously distributed (Therville N et al, 2019). The importance of this heterogeneity in tumor progression is unknown nowadays. In addition to ECM-induced compressive stress, ECM also influences the tumor progression *via* epithelial-mesenchymal transition (EMT) and further contributes to resistance to chemotherapeutic agents (Rice AJ et al, 2017).

Others have trialed therapeutic strategies that rely on decompression of solid tumor by reducing surrounding fibrosis and facilitating the accession of chemotherapeutic agents to the tumor (Lampi MC & Reinhart-King CA, 2018, Mohammadi H & Sahai E, 2018, Sheridan C, 2019). Some of these trials were stopped because they could accelerate the disease (Mohammadi H & Sahai E, 2018). As compressive forces decrease proliferation of most cancer cells in *in vitro* 3D models and here even induces cell death, by reducing the surrounding fibrosis, those therapeutics could allow an increase of cancer cell proliferation, survival and could promote tissue invasion and disease progression. These therapies should thus be considered in combination with conventional chemotherapies that can target newly proliferating cells freed from compressive context. It is even more important that compression of cancer cells without confinement increased cancer cell proliferative behaviour (Mary G et al, 2022). For these reasons, we are convinced that the balance between therapies reducing fibrosis and fibrosis-induced compression should be well mastered and individually adapted. Our findings also demonstrate that those mechanotherapeutics (modulating the mechanical environment in tumors) could also be used as a way to sensitize to oncogene targeted therapies such as PI3K inhibitors (Mohammadi H & Sahai E, 2018).

## 5. Material and Methods

### PI3K**α** genetic knockdown

MCF-7, MDA-MB-231, PANC-1 and CAPAN-1 cells were stably infected using lentiviral transduction containing p110α sh-RNA or scramble-sh-RNA into pLVTHM plasmid as described in (Thibault B et al, 2021), listed in Supporting Table 1 and selected positively by cell sorted flow cytometry using GFP expression.

### Cell culture

MCF-7 (ATCC; #CRL-3435), MDA-MB-231 (ATCC; # CRM-HTB-26), MDA-MB-468 (ATCC; # CRM-HTB-132), PANC-1 (ATCC; #CRL-1469), CAPAN-1 cells (ATCC; #HTB-79) and Mia-Paca-2 (ATCC; # CRM-CRL-1420) were cultured in Dulbecco’s Modified Eagle Medium (GIBCO; #61965026) supplemented with 10% fetal bovine serum (Eurobio Scientific; #CVFSVF00-01), Penicillin-Streptomycin 50 U/mL and 50 µg/mL respectively (Sigma-Aldrich; #P0781), L-Glutamine 2 mM (Sigma-Aldrich; #G7513) and Plasmocin 25 µg/mL (InvivoGen; #ant-mpp) and maintained at 37°C in humidified atmosphere with 5% CO2. Cells were tested for their absence of mycoplasma infection prior use. MCF-7 contains a constitutive PI3Kα activation (*PIK3CA*^E545K^) as compared to MDA-MB-231 cell line that harbors PI3Kα wild-type catalytic domain (*PIK3CA^WT^*) (Dunn S et al, 2019). Both pancreatic cancer cell lines present a constitutive PI3K activation (Thibault B et al, 2021).

### 2D compression

2×10^5^ MCF-7, MDA-MB-231, MDA-MB-468, PANC-1,CAPAN-1 or Mia-Paca-2 cells were plated in a 35 mm petri dish. MCF-7 and MDA-MB-231 were plated using Dulbecco’s Modified Eagle Medium (GIBCO; #61965026) supplemented with indicated products in previous paragraph, however without phenol red. 24h after plating, a 2% low gelling temperature agarose pad (Sigma-Aldrich; #A0701-25G) in Phosphate Buffered Saline (PBS) containing 0.5 mM MgCl_2_ and 0.9 mM CaCl_2_ respectively (Sigma-Aldrich; #D8662) was deposited on cells. 2D compression was calibrated at 200 Pa per cell adjusting the weight with small metal plates on 2% agarose pad and calculated using the formula: P=(1/CN)*(1/CS)*m*g (with P=Pressure in Pa; CN=Cell number; CS=Cell surface in m²; m=mass of the pad and metal plate in kg; g=gravitational force=9.80665 m/s-2). Controls were performed adding the volume of PBS 0.5 mM MgCl_2_ and 0.9 mM CaCl_2_ equivalent to the volume of the agarose pad. Each compression was performed for 24h, cells were harvested, and RNA/proteins were extracted.

### Normalized cell number assay

Cells were compressed following the protocol described above and/or treated with 10µM pan-PI3K inhibitor (GDC-0941; Axon Medchem) and/or Chloroquine (#C6628; Sigma-Aldrich) for 0h, 24h, 48h or 72h. After compression ± treatments, cells were rinsed with PBS (Eurobio Scientific, #CS1PBS 0101) and fixed for 15min using PBS containing 10% methanol and 10% acetic acid. Cells were stained for 15min using crystal violet (Sigma Aldrich; #HT90132). Images were performed using Chemidoc™ Imaging System (BioRad) and quantified using ImageJ software.

### Protein extraction

Cells were rinsed with PBS and detached by scrapping in PBS. Cells were harvested through a 5min centrifugation at 3,000g and 4°C. Cells were resuspended in a lysis buffer containing 150 mM NaCl, 50 mM Tris, 1 mM EDTA, 1% Triton, 2 mM dithiothreitol, 2 mM Sodium fluoride, 4 mM Sodium Orthovanadate and supplemented with protease inhibitors (Complete protease inhibitors, Roche). After a 20min incubation on ice, a 10min centrifugation was performed at 12,000g and 4°C and the supernatant was collected. Protein concentration was measured using Bicinchoninic assay (BC Assay Protein Quantification Kit, #3082, Interchim).

### Western Blot

20 µg of proteins were separated on a 10% polyacrylamide gel and transferred on nitrocellulose membranes (Amersham™ Protran®; #106000004) using Trans-Blot® Turbo™ (BioRad). Blocking was performed through a 1h incubation in 5% low fat milk in TBST. Membranes were incubated overnight at 4°C with primary antibodies in TBS-Tween 0.1%, 5% Bovine Serum Albumin as indicated in Supplementary Table 2. Then, membranes were rinsed three times with Tris-buffered saline 0,1% Tween (TBS-Tween 0.1%), incubated for 1h with secondary antibodies coupled to a horseradish peroxidase, in 1% low fat milk described in Supplementary Table 2. Membranes were next washed three times (1min, 5min and 10min) with TBS-Tween 0.1%. Proteins were detected through chemiluminescence (Clarity™ Western ECL Substrate; BioRad; #1705061) using Chemidoc™ Imaging System (BioRad) and quantified using ImageJ software. ≥3 independent replicates were performed for each protein analyzed.

### Immunostaining

Cells were cultured on glass coverslips. After compression, cells were fixed with 4% PFA in PBS for 10min and then permeabilized with 0.1% Triton in PBS for 5min. Cells were blocked in blocking solution (1% BSA in PBS) for 30min. Samples were incubated with p62/SQSTM1 primary antibody (Supplementary Table 2) diluted into blocking solution overnight at 4°C. Cells were washed with PBS and then incubated with the Alexa Fluor® 488 secondary antibody diluted into blocking solution for 1h (Supplementary Table 3). Samples were washed with PBS and incubated with DAPI (Sigma; #D9542 0.1µg/ml) as nuclear counterstain, for 3min. Coverslips were mounted in Fluoromount-G (Invitrogen; # 00-4958-02).

Images were acquired with a Plan Aprochromat 63x ON 1.4 oil immersion objective using a Zeiss LSM780 confocal Microscope.

### Annexin-V-FITC/Propidium Iodide (PI) assay

2×10^5^ MCF-7, MDA-MB-231, PANC-1 or CAPAN-1 cells were plated in a 35 mm petri dish. 2D compression was calibrated at 200 Pa per cell adjusting the weight with small metal plates on 2% agarose pad according to 2D compression paragraph and were treated with 10 µM GDC-0941 (pan-PI3K inhibitor; Axon Medchem) for 24h.

After 24h of compression, cells were washed with fresh phosphate-buffered saline (PBS), trypsinized (supernatants were kept) and stained with a FITC-Annexin-V/PI apoptosis detection kit (BD Pharmigen #556547) according to the manufacturer’s protocol. FITC-Annexin-V staining and PI incorporation were measured in cells with MACSQuant VYB Flow Cytometer and analyzed using FlowJo software.

### RNA extraction

Cells were rinsed twice with PBS. Cell lysis was performed by scraping cells in 1 volume of TRIzol reagent (Ambion; #15596018), and incubating 5min on ice. Then, 1/5 volume of chloroform (Sigma Aldrich; #32211) was added to the mix Trizol/cell lysate. After 3min incubation at room temperature, cell lysate was centrifuged at 12,000g and 4°C, for 10min and the supernatant was collected. RNA was precipitated by adding 1 volume of isopropanol (Sigma Aldrich; #33539), incubated for 10min at room temperature and then centrifuged at 12,000g and 4°C for 10min. The RNA pellet was rinsed using 70° ethanol. After a 5min centrifugation at 7,500g and 4°C the supernatant was removed, the RNA pellet was air dried for 5min and resuspended in 30µL of RNAse free water. RNA concentration was measured using NanoDrop™ 2000 (ThermoFischer scientific).

### Reverse transcription and quantitative PCR

cDNA synthesis was performed with iScript kit (BioRad; #1708891) according to the manufacturer protocol using C1000 Touch Thermal Cycler (BioRad) and the following conditions: annealing: 5min at 25°C, reverse transcription: 20min at 46°C, reverse transcriptase inactivation: 1min at 95°C. cDNAs were diluted to a half in RNAse free water. Gene expression was quantified with the SsoFast EvaGreen supermix kit (BioRad, #1725204) using a thermocycler (StepOne™ #4376374; Software v2.2.2) with the following conditions: 20sec at 95°C, 40 denaturation cycles: 3sec at 95°C, annealing and elongation: 30sec at 60°C. β-actin was used as housekeeping gene. The primers used are described in Supplementary Table 1 and each amplicon was validated by sequencing. Gene expression quantification was performed using the Livak method: 2^-(Delta^ ^Delta^ ^C(T))^ (Livak KJ & Schmittgen TD, 2001). Distinct RNA samples ≥4 were analyzed, each amplification was performed in technical duplicate.

### Transcriptomic analysis

Differential expression analysis under compressive stress in MCF-7 and MDA-MB-231 cells were performed from data provided by Kim *et al*., 2019 (Kim BG et al, 2019). MCF-7 and MDA-MB-231 breast cancer cells were gradually compressed (0; 0.3 to 0.8; 1.5 to 4.0; 7.0 to 8.0 kPa) and gene expression was quantified using Agilent-039494 SurePrint G3 Human GE v2 8×60K Microarray 039381, data available at https://www.ncbi.nlm.nih.gov/geo/query/acc.cgi?acc=GSE133134. The two PI3Kα and PI3Kβ inhibition signatures were performed from differential expression analysis after PI3Kα inhibition with BYL-719 (1 µM for 16 to 48 hours) in MCF-7 breast cancer cells (https://www.ncbi.nlm.nih.gov/geo/query/acc.cgi?acc=GSE64033) (Bosch A et al, 2015) or PI3Kβ inhibition after AZD8186 treatment (100mg/kg twice daily for 5 days) in HCC70 cells (https://www.ebi.ac.uk/arrayexpress/experiments/E-MTAB-4656/) (Lynch JT et al, 2017). For compressive stress and PI3Kα and PI3Kβ inhibition signatures, differential expression analysis was performed using DESeq2 v1.26.0 (Anders S & Huber W, 2010) on R version 3.6.3 with adjusted p-value <0.05. All differential expression data are relative data from normalized transcript per million (TPM). P-values on graphs were calculated using t-test comparing no compression condition (0 kPa) to each compression condition ([0.3-0.8], [1.5-4.0] or [7.0-8.0] kPa). Enrichment analysis were performed using Gene Set Enrichment Analysis software (GSEA), p-value <0.05 and Fold Discovery Rate (FDR)<25%.

### Statistical analysis

Comparisons between two experimental groups were performed using a paired two-tailed Student t-test. Comparisons between more than two experimental groups were performed using two-ways ANOVA. p-value<0.05 was considered significant. All analysis were performed using GraphPadPrism 9.3.1 software.

## Supporting information

Supplemental Figures for reviewer

## Data Availability

For this work, we utilized transcriptomic data from compressed breast cancer cells from Kim *et al*., 2019 (Kim BG et al, 2019), from Gene Expression Omnibus GEO accession number: GSE133134. Our work correlates compression data to PI3Kα inhibition data from Bosch *et al*., 2015 (Bosch A et al, 2015) GEO accession number: GSE64033) and PI3Kβ inhibition data from Lynch *et al*., 2017 (Lynch JT et al, 2017) ArrayExpress: ENA-ERP015852).

## Acknowledgments

We thank our colleagues for their critical reading of the manuscript. CS is mentored by Ecole Normale Supérieure de Lyon, Université Claude Bernard Lyon 1, France. We thank the Pôle technologique du Centre de Recherche en Cancérologie de Toulouse, for the technical support in imaging and cytometry platforms.

## Author Contributions

Experimentation, analysis, visualisation and interpretation of data: MDL, CS, EB, NT, MC, RDA, BT and JGG. Supervision of students: MDL, JGG. Methodology: MDL, NT, RDA, BT, MD and JGG. Drafting of the manuscript: MDL, CS and JGG. Revising the manuscript: MDL, BT, MD and JGG. Conceptualization of funded project, obtained funding, project administration, project supervision, validation: MD and JGG.

## 6. Figure legends

**Figure S1. Cell number in MCF-7/CAPAN-1 mechanically responsive and PANC-1/MDA-MB-231 non-responsive cells after 200 Pa compression and inhibition of class I PI3K. A.** Normalized cell number pictures of representative experiments performed in 6-well plates after 72h compression were shown in MCF-7, CAPAN-1, MDA-MB-231 and PANC-1 cells, treated with class I pan-PI3K inhibitor (GDC-0941 (10µM)) compared to control cells (CTL). **B.** Cell number pictures of representative experiments performed in 6-well plates after 72h compression were shown in PI3Kα genetic knockdown (sh PI3Kα) compared to control cells (CTL).

**Figure S2. Cell number and death events in mechanically responsive and non-responsive Mia-Paca-2/MDA-MB-468 pancreatic and breast cancer cells with inhibition of class I PI3K.** Normalized cell number was analyzed at 0, 24, 48 and 72 hours in Mia-Paca-2 and MDA-MB-468 and PANC-1, treated with class I pan-PI3K inhibitor (GDC-0941 (10µM)) compared to vehicle (CTL) (**A)** and after 200 Pa compression (200 Pa) and class I pan-PI3K inhibitor (200 Pa + GDC-0941 (10µM)) compared to 200 Pa compressed cells (200 Pa) **(B)**. After cell treatment, cell number was quantified using Crystal Violet staining. Results are presented as mean, +/- SEM, n=3. *p-value<0.05, **p-value<0.01. **C.** Representative western blots of p-AKT^S473^, AKT and GABARAP in mechanically responsive cells (Mia-Paca-2) and in mechanically non-responsive cells (MDA-MB-468) after 200 Pa compression (200 Pa) and non-compressed cells (0 Pa) for 24h. β-ACTIN was used as loading control. p-AKT^S473^/AKT, GABARAP and ACTIN quantitative analyses were performed using ImageJ software. Results are presented as mean, +/- SEM, n=4. *p-value<0.05.

**Figure S3. *In vitro* induction of cell death in MCF-7/CAPAN-1 mechanically responsive and PANC-1/MDA-MB-231 non-responsive cells after pharmaceutical inhibition of class I PI3K and 200 Pa compression.** After pan-PI3K inhibitor treatment and/or compression, cells were stained with a Annexin-V-FITC/PI apoptosis detection kit. Annexin-V-FITC staining and Propidium Iodide (PI) incorporation were measured in cells using a MACSQuant VYB Flow Cytometer and analyzed using FlowJo software. Early apoptotic cells correspond to the Annexin V positive and PI negative population. Late apoptotic and necrosis cells correspond to the Annexin V positive and PI positive population. Early and late apoptotic/necrosis cells populations were measured in MCF-7 (A), MDA-MB-231 (B), PANC-1 (C) and CAPAN-1 (D) cells in control (CTL), 200 Pa compressed (200 Pa), pan-PI3K inhibitor (GDC-0941 (10µM)) and 200 Pa compressed with pan-PI3K inhibitor (200 Pa + GDC-0941 (10µM)) conditions. Most representative n of n=4.

**Figure S4. PI3K signaling in MCF-7/CAPAN-1 mechanically responsive and PANC-1/MDA-MB-231 non-responsive cells after 200 Pa compression.** Representation of *PIK3CA, PIK3CB, PIK3CD, PIK3CG* mRNA expressions in mechanically responsive cells (MCF-7/CAPAN-1) and in mechanically non-responsive cells (PANC-1/MDA-MB-231) after 200 Pa compression (200 Pa) and non-compressed cells (0 Pa) for 24h. β-ACTIN was used as housekeeping gene. Results are presented as mean, +/- SEM. n≥4 for each gene. n=4. p=p-value. *p-value<0.05.

**Figure S5. YAP/TAZ, PI3K-AKT and autophagy pathways under increasing compressive stress in breast cancer cells.** MDA-MB-231 (PIK3CA^wt^; blue triangles) and MCF-7 (PIK3CA^E545K^ mutated, black squares) breast cancer cells were gradually compressed (0 kPa; [0.3 to 0.8 kPa]; [1.5 to 4.0 kPa]; [7.0 to 8.0 kPa] respectively) and gene expression was quantified using Agilent microarray (Data available in https://www.ncbi.nlm.nih.gov/geo/query/acc.cgi?acc=GSE133134). Differentially expressed genes (p-value<0.05) were correlated. **A.** *ACTINB* (1 gene), *LAMINA1* (1 gene) and *LAMIN* gene class (12 genes) expressions were represented as housekeeping genes. **B.** Regulation of REACTOME PI3K-AKT signature (102 genes) and *PI3KCA*, *PIK3CB* and *PIK3CD* differential gene expressions were represented (± SEM. *<0.05; ***<0.001). **C.** Regulation of YAP/TAZ *via* HIPPO and non HIPPO (45 genes) and autophagy (35 genes) pathways were represented in breast cancer cells (MCF-7 and MDA-MB-231) gradually compressed. ± SEM. p: *<0.05; **<0.01; ***<0.001.

**Figure S6. YAP/TAZ pathway in MCF-7/CAPAN-1 mechanically responsive and PANC-1/MDA-MB-231 non-responsive cells after 200 Pa compression. A)** Mechanically responsive cells (MCF-7/CAPAN-1) and in mechanically non-responsive cells (PANC-1/MDA-MB-231) were 200 Pa compressed for 24h and *TEAD1*, *c-JUN*, *CCN1* and *CCN2* mRNA expressions were quantified. β-ACTIN was used as housekeeping gene. **B)** Mechanically responsive cells (MCF-7/CAPAN-1) and in mechanically non-responsive cells (PANC-1/MDA-MB-231) were 200 Pa compressed for 24h and p-YAP^S127^/YAP protein expressions were quantified. β-ACTIN was used as loading control. n≥4. p=p-value. *p-value<0.05.

