## Supplementary figures and images for "Mechanical compressive forces increase PI3K output signaling in breast and pancreatic cancer cells"

### Supplemental Figures for reviewer

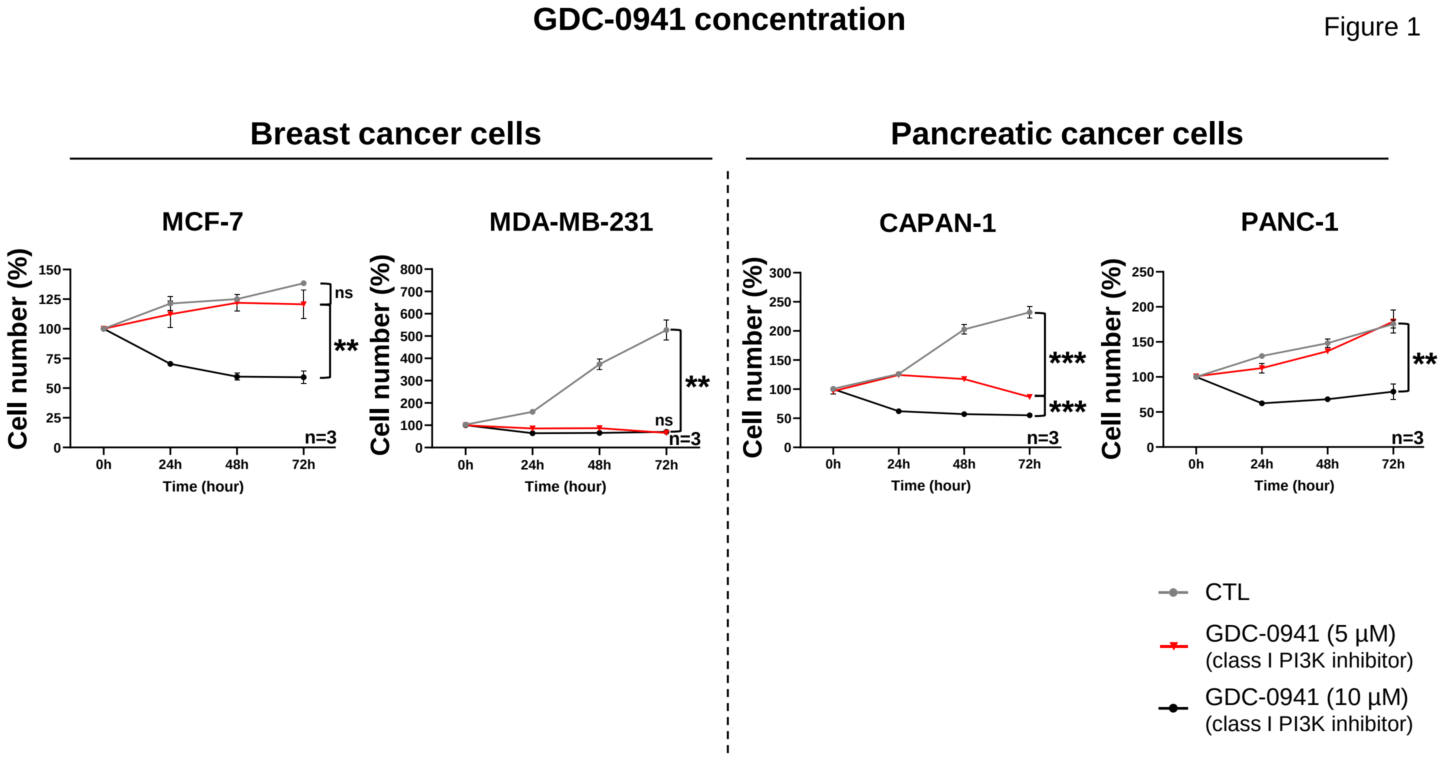

Figure 2

#### Nuclei sizes

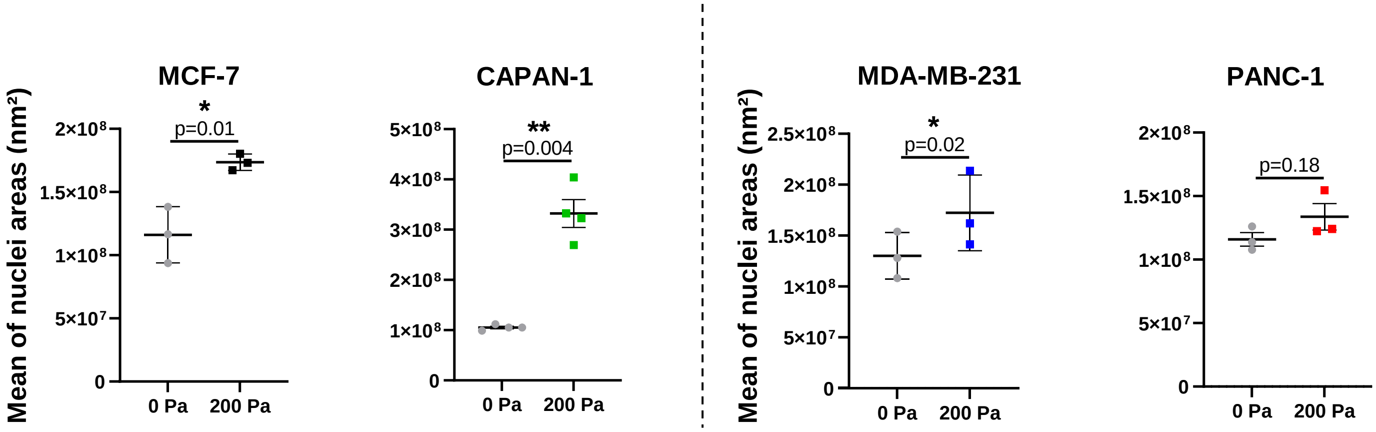

Figure 3

#### Hypoxia experiment

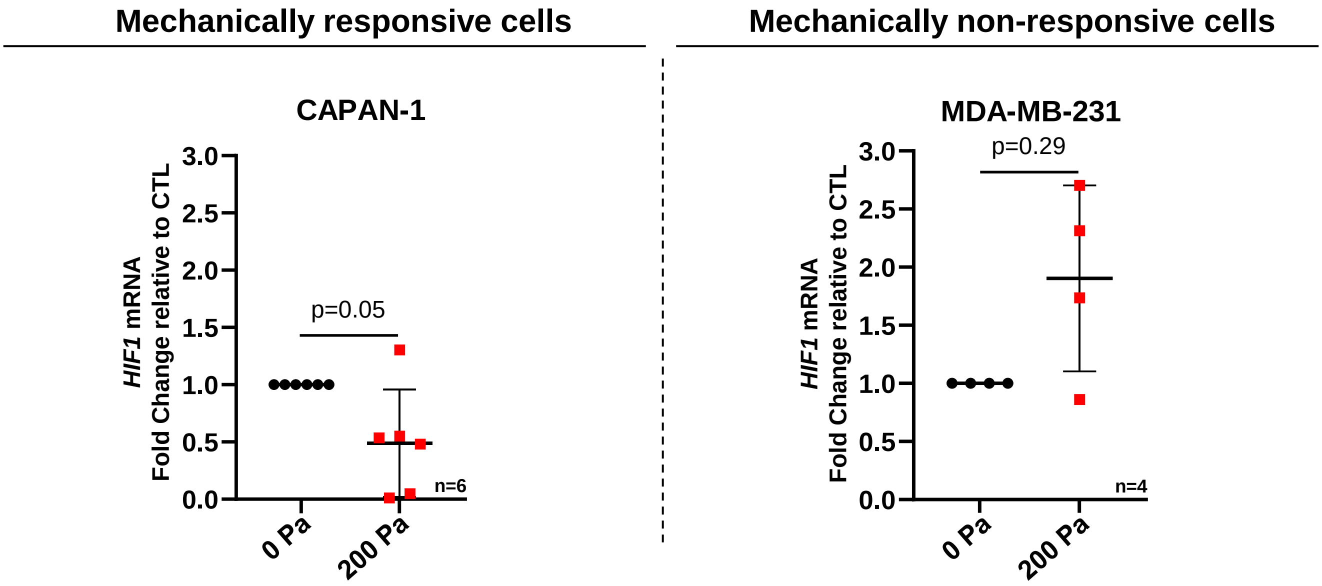

Figure 4

#### Compression with glass beads

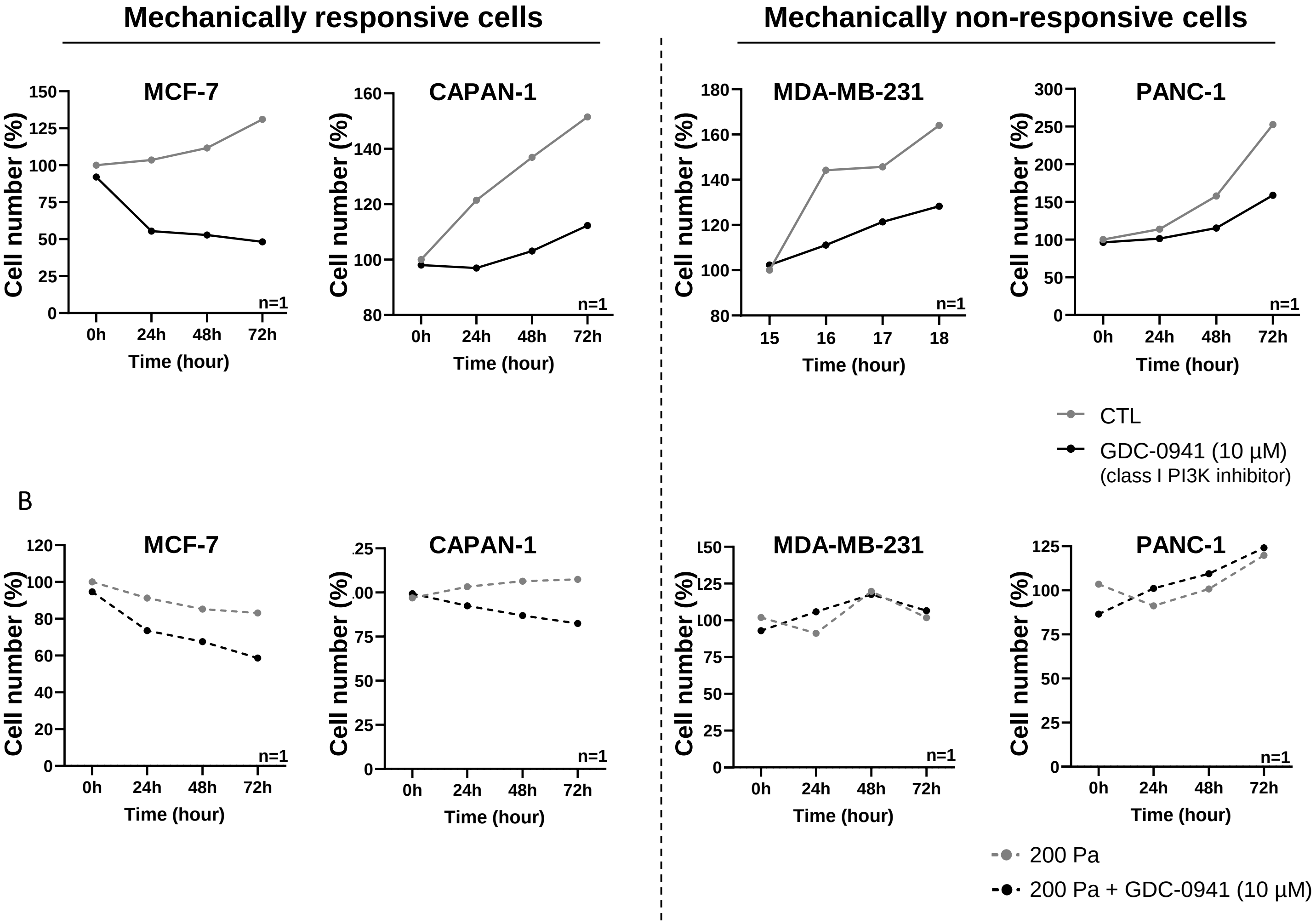

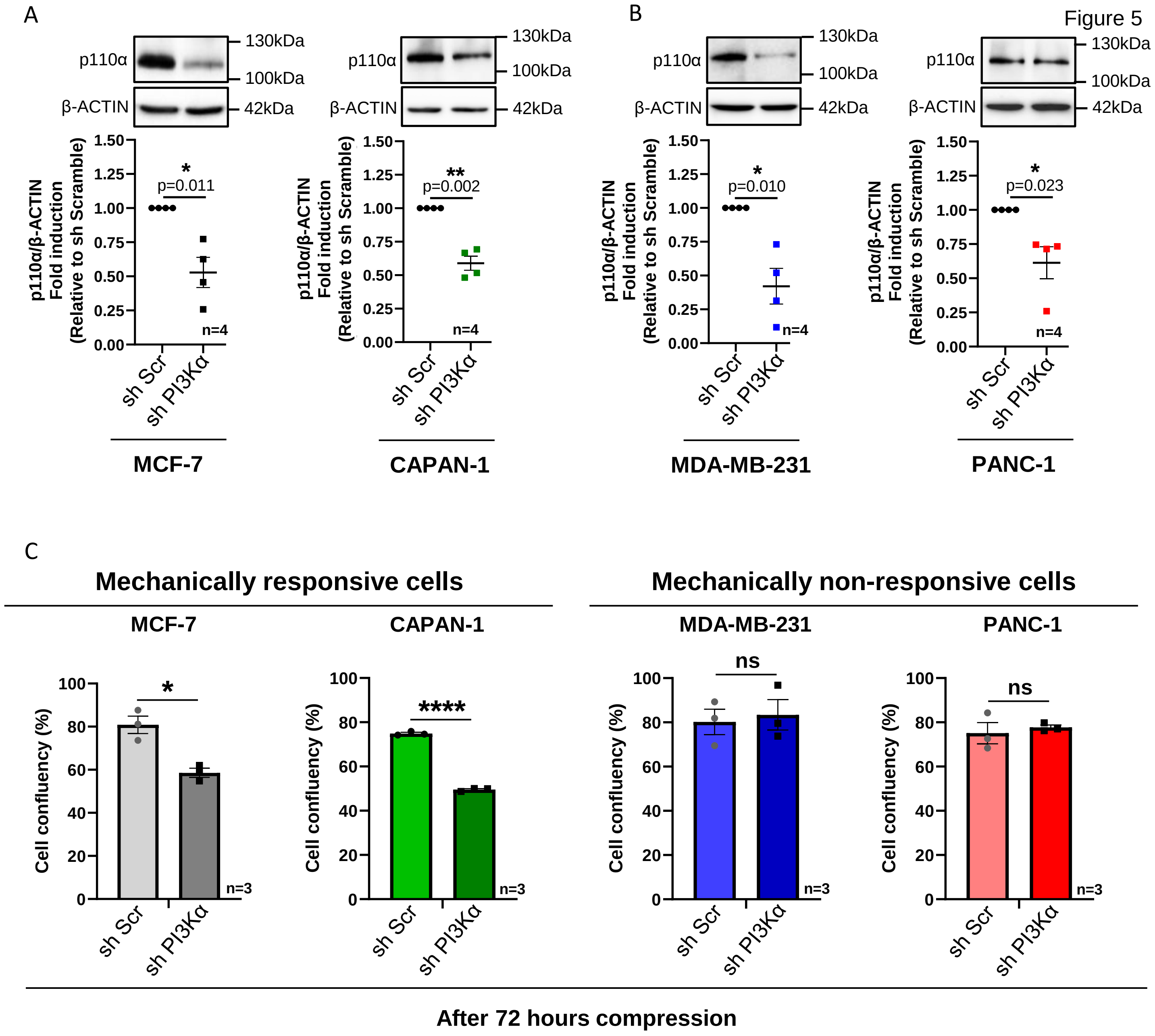
