## Supplemental Figures for "Mechanical compressive forces increase PI3K output signaling in breast and pancreatic cancer cells"

Figure S1

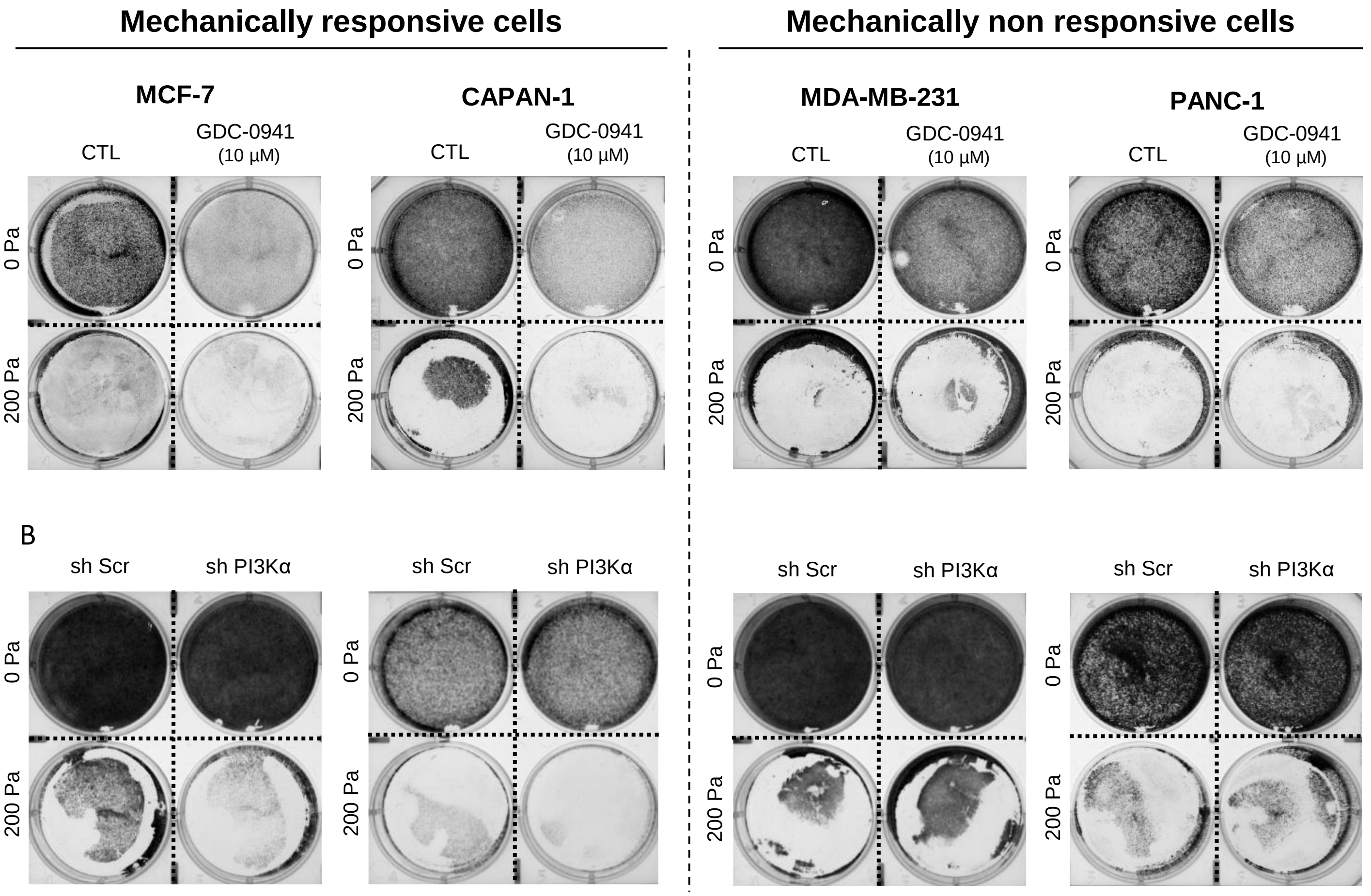

Mechanically responsive cells
Mechanically non responsive cells
MCF-7
CAPAN-1
MDA-MB-231
PANC-1
CTL
GDC-0941 (10 µM)
0 Pa
200 Pa
B
sh Scr
sh PI3Kα

Figure S2

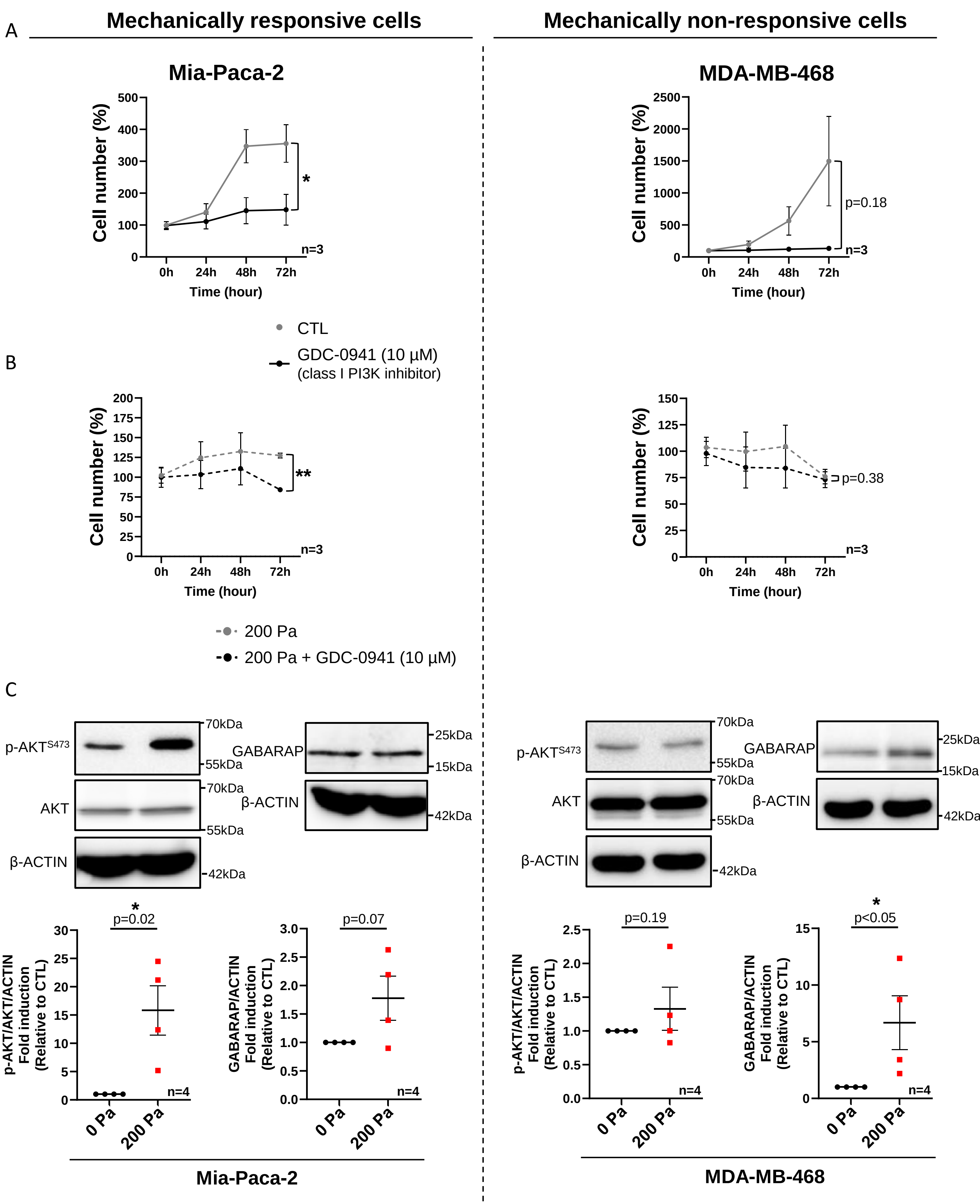
A
Mechanically responsive cells
Mechanically non-responsive cells
Mia-Paca-2
Cell number (%)
500
400
300
200
100
0
0h
24h
48h
72h
Time (hour)
*
n=3
MDA-MB-468
Cell number (%)
2500
2000
1500
1000
500
0
0h
24h
48h
72h
Time (hour)
p=0.18
n=3
CTL
GDC-0941 (10 µM)
(class I PI3K inhibitor)
B
Cell number (%)
200
175
150
125
100
75
50
25
0
0h
24h
48h
72h
Time (hour)
**
n=3
Cell number (%)
150
125
100
75
50
25
0
0h
24h
48h
72h
Time (hour)
p=0.38
n=3
200 Pa
200 Pa + GDC-0941 (10 µM)
C
p-AKT^S473
70kDa
55kDa
AKT
70kDa
55kDa
β-ACTIN
42kDa
GABARAP
25kDa
15kDa
β-ACTIN
42kDa
p-AKT^S473
70kDa
55kDa
AKT
70kDa
55kDa
β-ACTIN
42kDa
GABARAP
25kDa
15kDa
β-ACTIN
42kDa
p-AKT/AKT/ACTIN Fold induction (Relative to CTL)
*
p=0.02
30
25
20
15
10
5
0
n=4
0 Pa
200 Pa
GABARAP/ACTIN Fold induction (Relative to CTL)
p=0.07
3.0
2.5
2.0
1.5
1.0
0.5
0.0
n=4
0 Pa
200 Pa
p-AKT/AKT/ACTIN Fold induction (Relative to CTL)
p=0.19
2.5
2.0
1.5
1.0
0.5
0.0
n=4
0 Pa
200 Pa
GABARAP/ACTIN Fold induction (Relative to CTL)
*
p<0.05
15
10
5
0
n=4
0 Pa
200 Pa
Mia-Paca-2
MDA-MB-468

#### Mechanically responsive cells

##### MCF-7

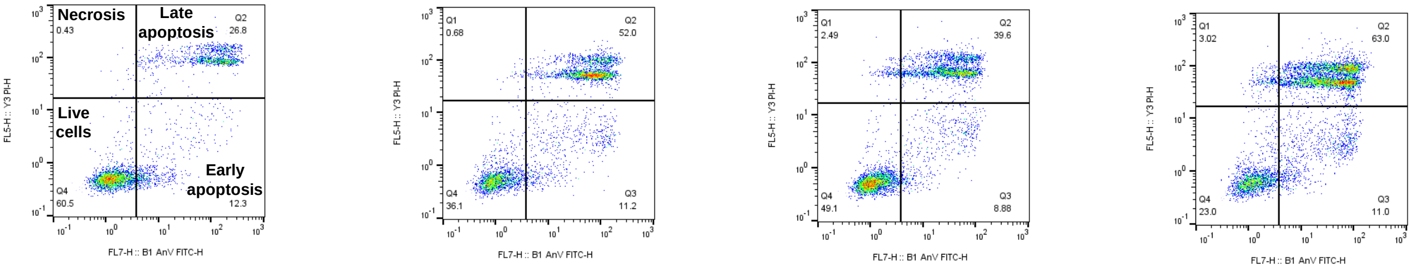


CTL | 200 Pa | GDC-0941 | 200 Pa + GDC-0941

Most representative n of n=4

##### CAPAN-1

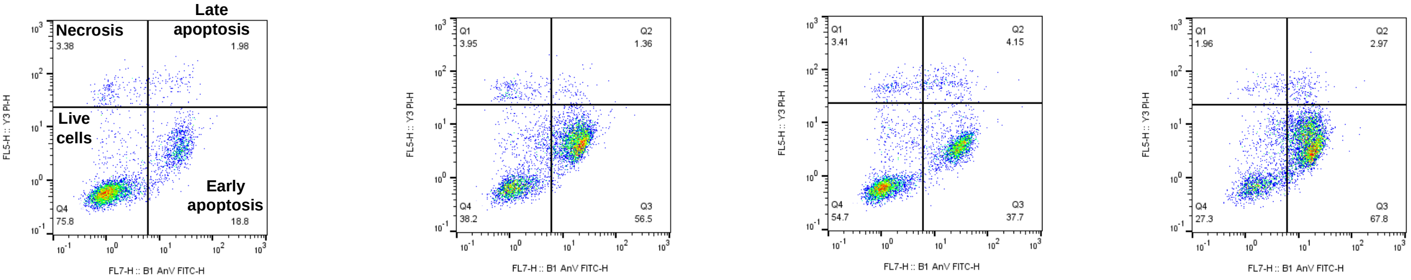


CTL | 200 Pa | GDC-0941 | 200 Pa + GDC-0941

Most representative n of n=4

#### Mechanically non responsive cells

##### MDA-MB-231

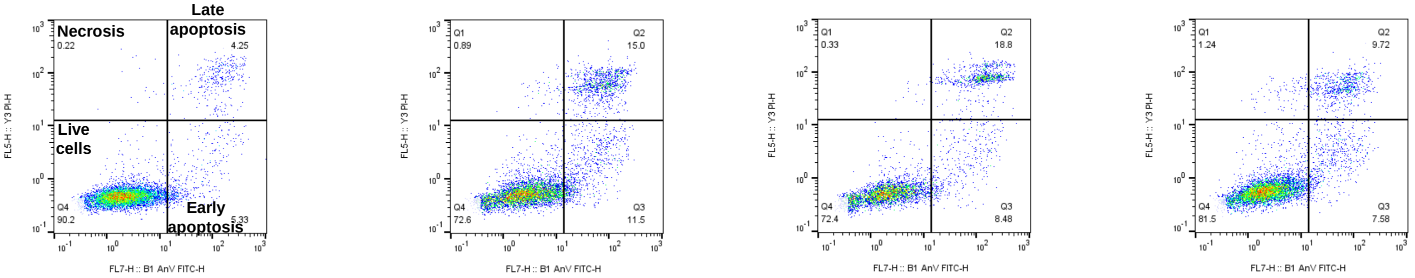


CTL | 200 Pa | GDC-0941 | 200 Pa + GDC-0941

Most representative n of n=4

##### PANC-1

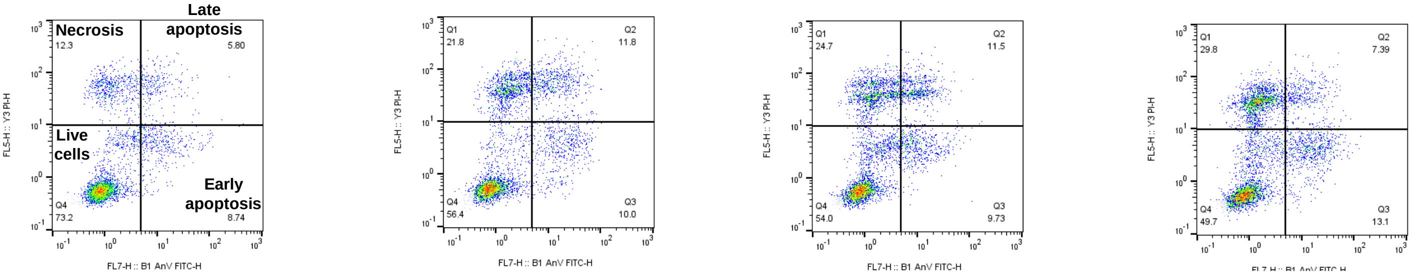


CTL | 200 Pa | GDC-0941 | 200 Pa + GDC-0941

Most representative n of n=4

Figure S4

### PI3K pathway

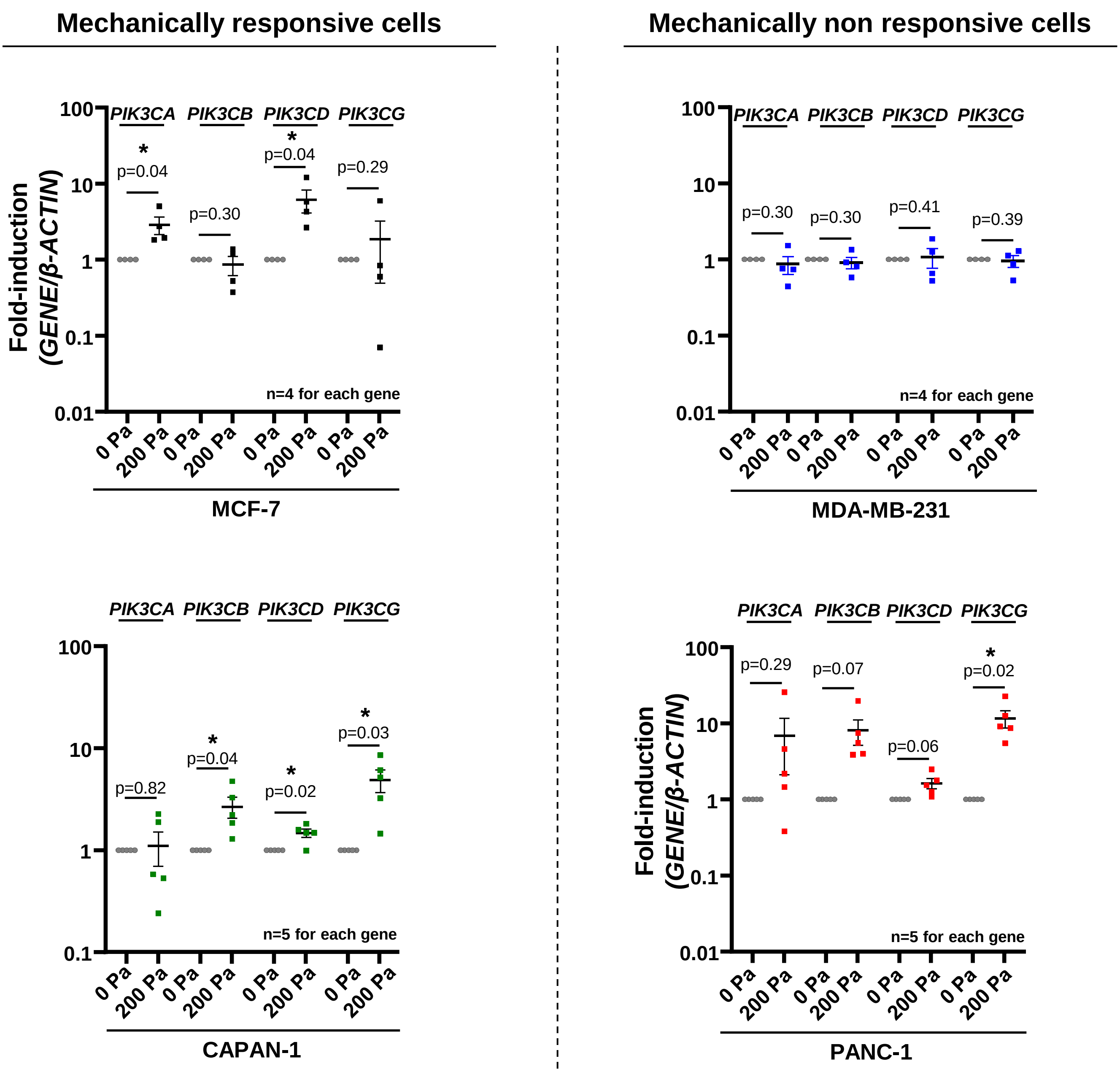

Figure S5

A

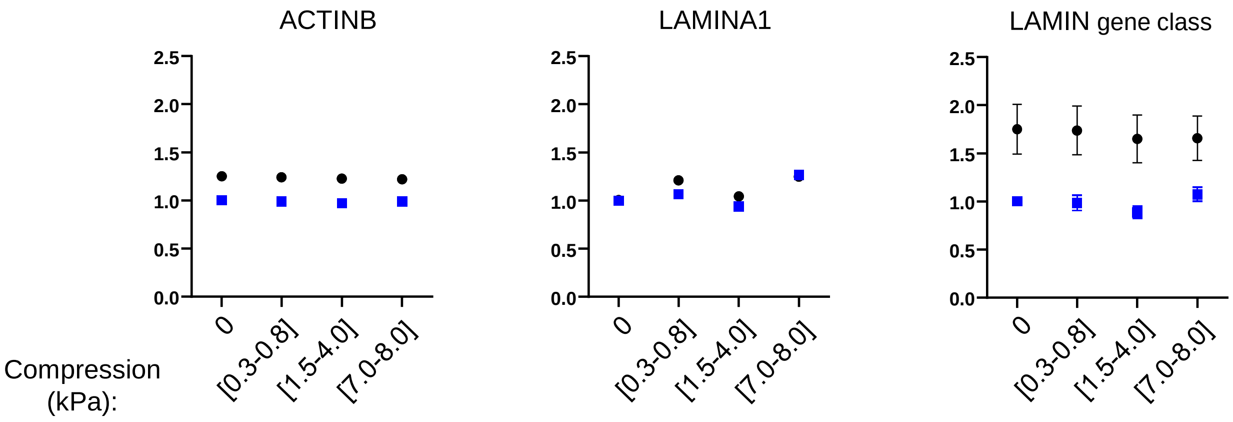


B

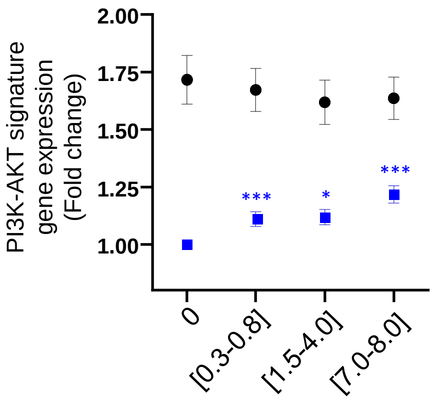


| | MCF-7 | | | | MDA-MB-231 | | | |
|---|---|---|---|---|---|---|---|---|
| Pressure (kPa) | [0] | [0.3-0.8] | [1.5-4.0] | [7.0-8.0] | [0] | [0.3-0.8] | [1.5-4.0] | [7.0-8.0] |
| *PIK3CA* | 1.00 | 0.96 | 0.97 | 1.06 | 1.00 | 0.91 | 0.91 | 0.92 |
| *PIK3CB* | 1.00 | 1.29 | 1.30 | 1.15 | 1.00 | 0.90 | 0.86 | 0.85 |
| *PIK3CD* | 1.00 | 1.07 | 0.99 | 2.11 | 1.00 | 0.76 | 0.91 | 0.77 |

C

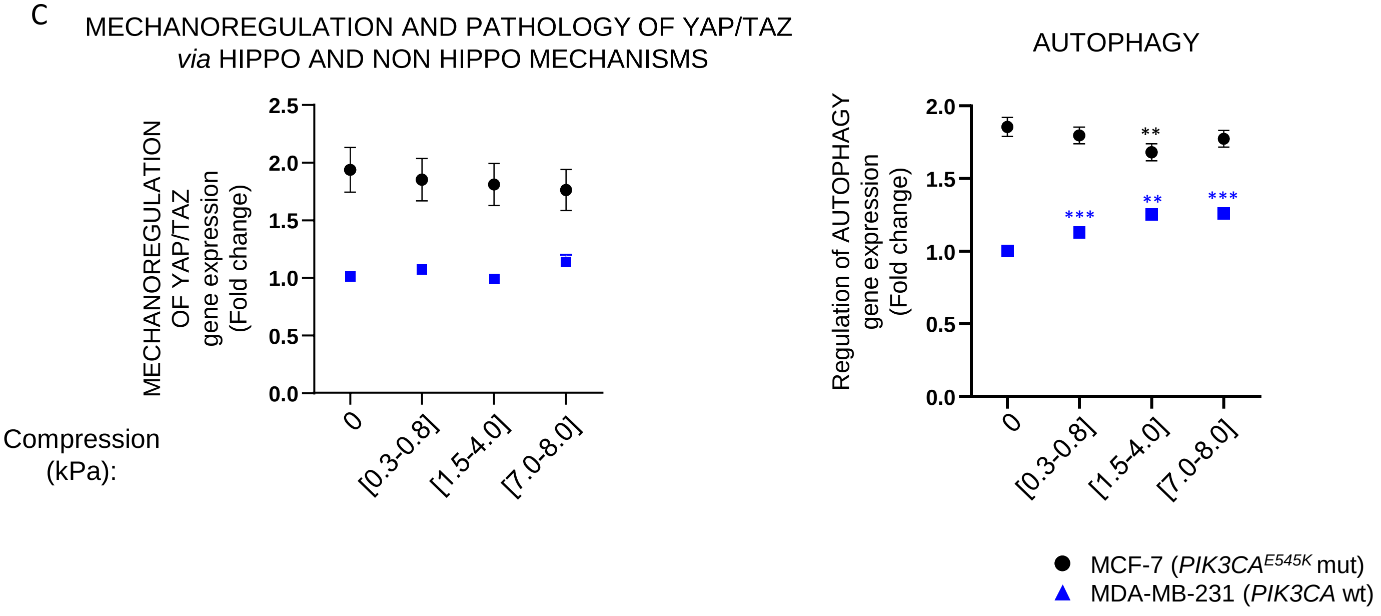

Figure S6

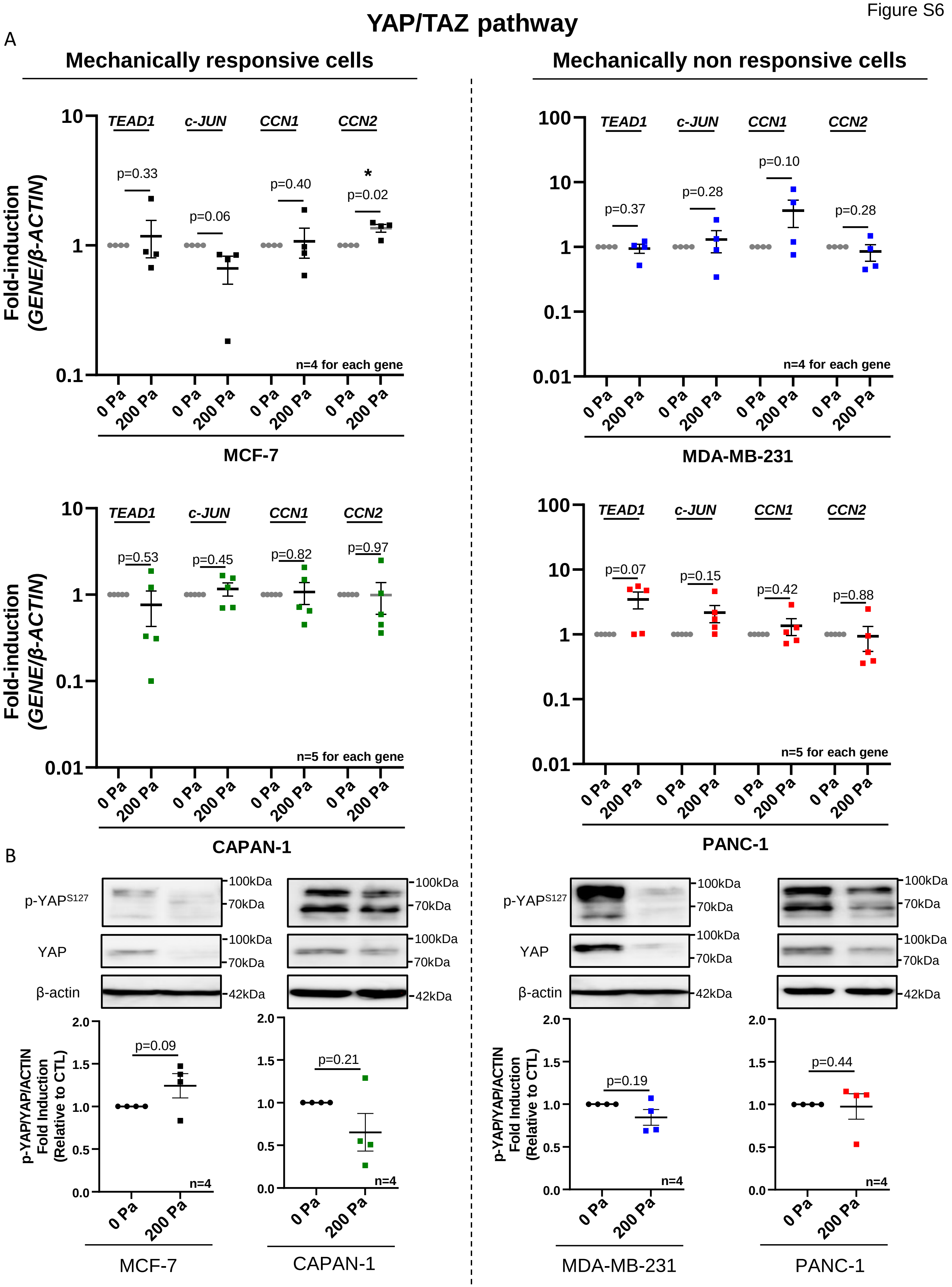
YAP/TAZ pathway
A
Mechanically responsive cells
Mechanically non responsive cells
Fold-induction (GENE/β-ACTIN)
TEAD1
c-JUN
CCN1
CCN2
p=0.33
p=0.06
p=0.40
*
p=0.02
n=4 for each gene
0 Pa
200 Pa
MCF-7
p=0.37
p=0.28
p=0.10
p=0.28
MDA-MB-231
p=0.53
p=0.45
p=0.82
p=0.97
n=5 for each gene
CAPAN-1
p=0.07
p=0.15
p=0.42
p=0.88
PANC-1
B
p-YAPS127
YAP
β-actin
100kDa
70kDa
42kDa
p-YAP/YAP/ACTIN Fold Induction (Relative to CTL)
p=0.09
p=0.21
p=0.19
p=0.44
n=4
MCF-7
CAPAN-1
MDA-MB-231
PANC-1
