## Supplemental Table for "Mechanical compressive forces increase PI3K output signaling in breast and pancreatic cancer cells"

### Supporting Tables

#### *Supporting Table 1. List of primers*

| APPLICATION | GENE | SEQUENCE |
|---|---|---|
| RT-qPCR | Homo Sapiens | |
| **Housekeeping gene** | *ACTB* | Forward: 5’-CTCCATCATGAAGTGTGACGTG-3'<br>Reverse: 5’-GGAGTACTTGCGCTCAGG-3' |
| **PI3K pathway** | *PIK3CA* | Forward: 5'-GTATCCCGAGAAGCAGGATTTAG -3'<br>Reverse: 5'-CAGAGAGAGGATCTCGTGTAGAA-3‘ |
| | *PIK3CB* | Forward: 5'-ATGGGTGAGCCTCTTCTTTATG -3'<br>Reverse: 5'-CCTATTCCTGAGGTTGGTTTGT-3' |
| | *PIK3CD* | Forward: 5'-CCCACAGGTGATCCTAACATATC-3'<br>Reverse: 5'-ACTTCTGGCTCTGTTGAGTTT-3' |
| | *PIK3CG* | Forward: 5'- GATTCTTCTTCCTTGCCCTTTG-3'<br>Reverse: 5'-CAGGTGGGTAGAGTGTGATTT-3' |
| **Autophagy pathway** | *GABARAPL1* | Forward: 5'-CCAGTACAAGGAGGACCATCC-3'<br>Reverse: 5'-GAATAAGGCGTCCTCAGGTCTC-3' |
| **YAP/TAZ pathway** | *TEAD1* | Forward: 5’-CAGGAGGAGACTCTCCCTG-3’<br>Reverse: 5’-CCTCCTGAAAGCTTTGCTCG-3’ |
| | *c-JUN* | Forward: 5'-GGTCGGCAGTATAGTCCGAAC-3'<br>Reverse: 5'-CTTTCCGCCGCTGTCAAC-3' |
| | *CCN2* | Forward: 5’-CCTATTCTGTCACTTCGGCTC-3’<br>Reverse: 5’-CAGACGAACGTCCATGCTG-3’ |
| | *CCN1* | Forward: 5'-GCTCTGAAGGGGATCTGC-3'<br>Reverse: 5'-GTAACTTTGACCAGCCGAGG-3' |
| **ShRNA** | Homo Sapiens | |
| | *Sh-RNA-Scramble* | Forward:<br>5′CGCGTCCCCTTCTAGAGATAGTCTGTACGTTTCAAGAGAACGTACAGACTATCTCTAGAA TTTTTGGAAAT-3’<br><br>Reverse:<br>5′-CGATTTCCAAAAATTCTAGAGATAGTCTGTACGTTCTCTTGAAACGTAC AGACTATCTCTAGAAGGGGA |
| | *Sh-RNA-p110α1* | Forward:<br>5’-CGCGTCCCCGCGAAATTCTCACACTATTATTTCAAGAGAATAATAGTGT GAGAATTTCGCTTTTTGGAAAT-3’<br>Reverse:<br>5'- CGATTTCCAAAAAGCGAAATTCTCACACTATTATTCTCTTGAAATAAT AGTGTGA-GAATTTCGCGGGGA |
| | *Sh-RNA-p110α2* | Forward:<br>5’-CGCGTCCCCGCACAATCCATGAACACATTTTCAAGGAAATGCTGTTCA TGGATTGTGCTTTTTGGAAAT<br>Reverse:<br>5'- CGATTTCCAAAAAGCACAATCCATGAACACATTTCTCTGAAAATGCTG TTCATGGATTGTGCGGGGA-3’ |

#### *Supporting Table 2. Primary antibodies*

| PRIMARY ANTIBODY | SPECIES | SOURCE | REFERENCE NUMBER | DILUTION IF/WB |
|---|---|---|---|---|
| β-ACTIN | Mouse | Sigma Aldrich | #A2228 | 1/10000 |
| AKT | Rabbit | Cell Signaling | #4691 | 1/1000 |
| GABARAP | Rabbit | Cell Signaling | #13733 | 1/1000 |
| LC3B | Rabbit | Cell Signaling | #2775 | 1/1000 |
| p110α | Rabbit | Cell Signaling | #4249 | 1/500 |
| p110β | Rabbit | Santa Cruz Biotechnology | #602 | 1/500 |
| p62/SQSTM1 | Rabbit | Cell Signaling | #7695 | IF: 1/500 |
| p-AKT(Ser473) | Rabbit | Cell Signaling | #4060 | 1/2000 |
| p-YAP(Ser127) | Rabbit | Cell Signaling | #4911 | 1/1000 |
| YAP | Rabbit | Cell Signaling | #4912 | 1/1000 |

#### *Supporting Table 3. Secondary antibodies*

| SECONDARY ANTIBODY | SPECIES | SOURCE | REFERENCE NUMBER | DILUTION WB |
|---|---|---|---|---|
| Anti-rabbit IgG-Horse Radish Peroxidase | Goat | Invitrogen | #31460 | 1/5000 |
| Anti-mouse IgG-Horse Radish Peroxidase | Goat | Invitrogen | #31430 | 1/10000 |
| Anti-rabbit Alexa Fluor 488 | Goat | Abcam | #ab150077 | 1/200 |
